# Extensive PFAS accumulation by human gut bacteria

**DOI:** 10.1101/2024.09.17.613493

**Authors:** Anna E. Lindell, Anne Grießhammer, Lena Michaelis, Dimitrios Papagiannidis, Hannah Ochner, Stephan Kamrad, Rui Guan, Sonja Blasche, Leandro Ventimiglia, Bini Ramachandran, Hilal Ozgur, Aleksej Zelezniak, Nonantzin Beristain-Covarrubias, Juan Carlos Yam-Puc, Indra Roux, Leon P. Barron, Alexandra K. Richardson, Maria Guerra Martin, Vladimir Benes, Nobuhiro Morone, James E. D. Thaventhiran, Tanmay A.M. Bharat, Mikhail Savitski, Lisa Maier, Kiran R. Patil

**Author notes:** Correspondence to: KRP.

## Abstract

Per- and polyfluoroalkyl Substances (PFAS) – the so-called ‘forever chemicals’ – are a major cause of environmental and health concern due to their toxicity and long-term persistence^1,2^. Yet, no efficient mechanisms for their removal have been identified. Here we report bioaccumulation of PFAS by several gut bacterial species over a wide range of concentrations from nanomolar up to 500 μM. For bioaccumulating Bacteroides uniformis, a highly prevalent species, we estimate intracellular PFAS concentration in the mM range – above that of most native metabolites. Despite this high bioaccumulation, B. uniformis cells could grow appreciably up to 250 μM perfluorononanoic acid (PFNA) exposure. Escherichia coli, which accumulated PFAS to a much lesser extent, substantially increased PFAS bioaccumulation when lacking TolC efflux pump indicating trans-membrane transport in PFAS bioaccumulation. Electron microscopy and cryogenic Focused Ion Beam-Secondary Ion Mass-spectrometry revealed distinct morphological changes and intracellular localisation of PFNA aggregates. Bioaccumulation of PFAS and transmembrane transport is also evident in proteomics, metabolomics, thermal proteome profiling, and mutations following adaptive laboratory evolution. In an in vivo context, mice colonized with human gut bacteria showed, compared to germ-free controls or those colonized with low-bioaccumulating bacteria, higher PFNA levels in excreted feces. As the gut microbiota is a critical interface between exposure and human body, our results have implications for understanding and utilizing microbial contribution to PFAS clearance.

---

The environmental contamination by manufactured chemicals has by some estimates exceeded the safe planetary boundary^3,4^. Due to the widespread contamination of water and agricultural systems, a vast number of chemical pollutants make it into the food and hence into the human body^1,2,5–7^. The gut microbiota is particularly susceptible to exposure and adverse interactions therein could cause systemic effects on the host due to the critical role of the microbiota in host physiology^8,9^. To assess the potential impact of food-borne pollutants on commensal gut bacteria, we screened a panel of 42 common pollutants against 14 representative gut bacterial strains. The bacteria were selected for their prevalence and abundance in healthy population and for phylogenetic and metabolic representation^10,11^ (SI Table 1). Pollutants were selected by considering reported occurrence in food and representation of different classes including pesticides, food contact materials and industrial chemicals (SI Table 2).

We started with a community-based screening approach, wherein we assessed the ability of a mix of gut bacterial strains to sequester the pollutant compounds during 4 h exposure (SI Fig. 1a). Thirteen pollutants were found to be depleted to more than 20 % by one or both synthetic communities (SI Fig. 1b). Ten compounds were then tested for depletion by 14 individual strains during a 24 h growth period (SI Fig. 1c); seven pollutants were found to be depleted to more than 20 % by at least one of the bacterial strains (Fig. 1a). To our knowledge, most of these interactions have not been reported before. By comparing the whole culture and supernatant concentrations to compound controls we were able to distinguish the observed sequestration between bioaccumulation and biotransformation. Bioaccumulation – storage of the pollutant without modification – was defined as compound sequestration to at least 20 % from the supernatant but recovery from the whole culture sample, whereas biotransformation was defined as both supernatant and whole culture sample showing more than 20 % depletion. These interactions provide a starting point to mechanistically understand the role of the microbiota in pollutant toxicokinetics.

**Figure 1.**
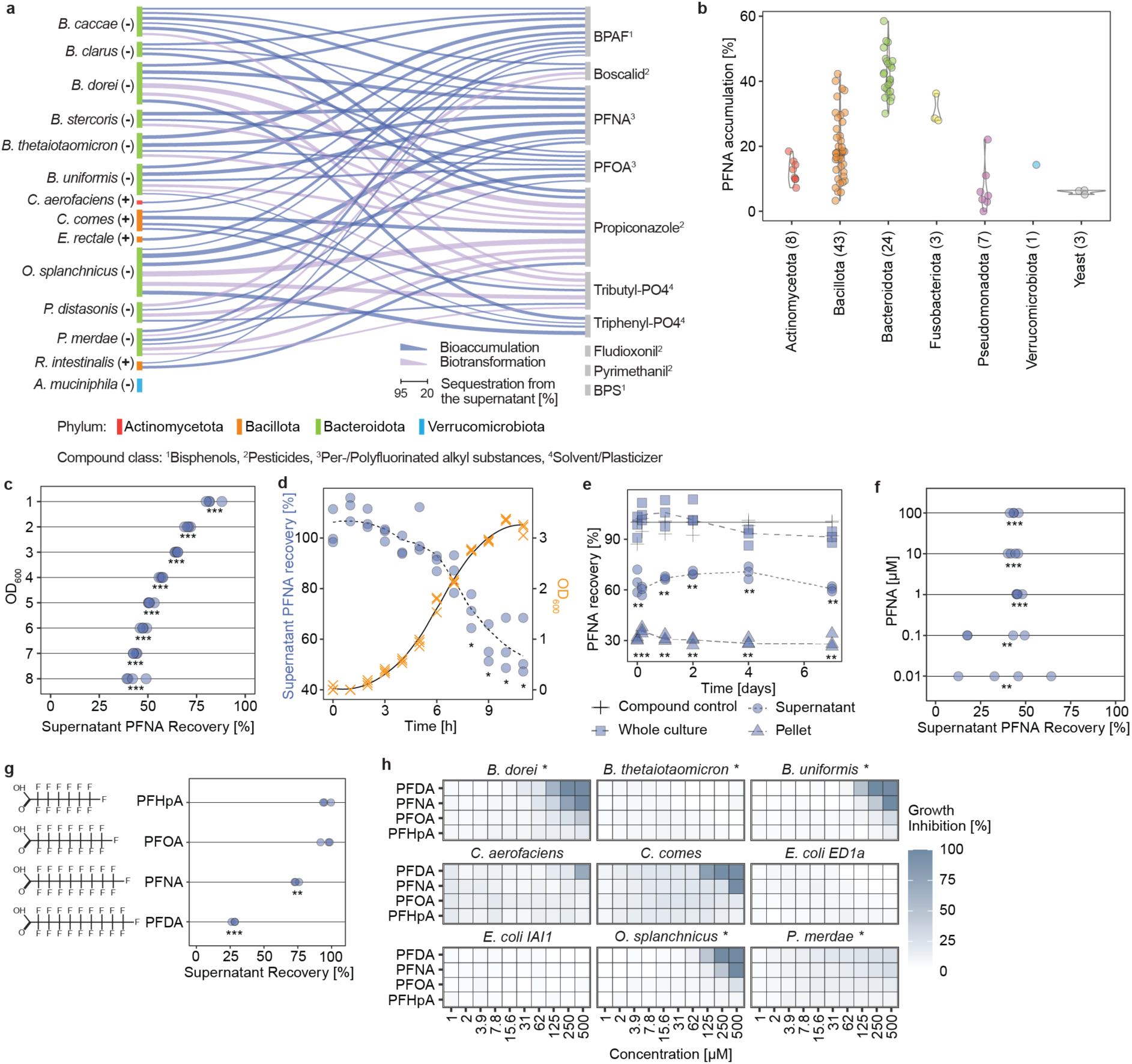
Abundant gut bacterial species bioaccumulate and biotransform common chemical pollutants and accumulate and tolerate PFAS over a broad concentration range. **a.** Specificity of human gut bacteria to sequester (bioaccumulate/biotransform) chemical pollutants during 24 h growth period as identified using mass-spectrometry. Links between bacterial species and pollutant denotes >20 % depletion. The link thickness is proportional to the median depletion from 6 replicates (3 biological, 2 technical, initial pollutant concentration = 20 μM) (SI Table 4,5). **b.** 89 strains were tested for PFNA accumulation. Selected strains cover all major bacterial phyla and three yeasts. Numbers in brackets show the number of strains from each phylum. Bacterial strains from the same phylum show similar PFNA accumulating capacity (Optical density (OD600) = 3.75; initial PFNA concentration = 20 μM (9,280 µg/l)); n = 3 technical replicates (SI Table 6,7). **c.** PFNA depletion by *B. uniformis* cultures of varying OD600 in PBS buffer (OD600 = 1-8; initial PFNA concentration = 20 μM (9,280 µg/l)). All ODs show significant PFNA accumulation compared to the compound control. *** p < 0.001. n = 4 technical replicates (SI Table 8). **d.** Kinetics of PFNA depletion during *B. uniformis* growth starting with low cell density (initial OD600 = 0.05; initial PFNA concentration = 20 μM (9,280 µg/l)). A significant amount of PFNA was sequestered from the media after 8 h of growth and onwards. * p<0.05 and >20% PFNA sequestration from the media compared to the compound control; n = 3 biological replicates (SI Table 9.10). **e.** Kinetics of PFNA depletion by *B. uniformis* when starting with a high cell density in PBS (OD600 = 3.75; initial PFNA concentration = 20 μM (9,280 µg/l)). Significant bioaccumulation of ca. 30 % PFNA, which is not released within 7 days. ** p-value < 0.01 (supernatant compared to the compound control; pellet compared to 0); n = 3 biological replicates (SI Table 11). **f.** PFNA is significantly bioaccumulated by *B. uniformis* grown in mGAM at a range of initial concentrations (initial OD600 = 0.05; initial PFNA concentrations = 0.01 to 100 μM (4.64 µg/l – 232 mg/l)) compared to the compound control. ** p-value < 0.01; *** p-value < 0.001; n = 4 technical replicates (SI Table 13). **g.** Bioaccumulation of PFAS compounds with varying chain length by *B. uniformis* (OD600 = 3.75; initial concentration for all compounds = 20 μM (PFHpA 7280 µg/l, PFOA 8,280 µg/l, PFNA 9,280 µg/l, PFDA 10,280 µg/l)). PFNA and PFDA are significantly accumulated compared to the compound control. ** p-value < 0.01; *** p-value < 0.001; n = 3 technical replicates (SI Table 14). **h.** Growth sensitivity of gut bacteria to PFAS is independent of bioaccumulation (n = 3 technical replicates); *Bioaccumulating bacteria (SI Table 16,17).

The pollutants bioaccumulated by gut bacteria included perfluorooctanoic acid (PFOA) and perfluorononanoic acid (PFNA), which belong to the chemical group of PFAS (per- and polyfluoroalkyl substances). This was surprising as no substantial microbial PFAS sequestration has been reported to our knowledge. Nine of the tested bacterial species – *Bacteroides caccae, B. clarus, B. dorei, B. stercoris, B. thetaiotaomicron, B. uniformis, Odoribacter splanchnicus, Parabacteroides distasonis, P. merdae* – bioaccumulated PFNA and/or PFOA. The degree of bioaccumulation over 24 h for 20 μM (9,280 µg/l) PFNA exposure varied from 25 % (*P. distasonis*) to 74 % (*O. splanchnicus*) and for 20 μM (8,280 µg/l) PFOA from 23 % (*P. merdae*) to 58 % (*O. splanchnicus*). PFAS are widely used in manufacturing and various consumer products due to their exceptional surfactant properties and stability. Yet, these same properties have made PFAS of great concern for environmental and human health^1,3,12–16^. The annual health-related cost of PFAS exposure has been estimated to be between 50 and 80 billion Euros across Europe^17^. These widespread concerns for PFAS have led to manufacturing/use limitations for several compounds^18,19^ and further legislative actions are being taken to either control PFAS levels in drinking water^20^ or ban PFAS manufacturing/use entirely^21^. Yet, such efforts are not global and with long environmental half-lives and no efficient route for their removal, PFAS pose an urgent challenge for both planetary and human health. Chemical methods to degrade PFAS have so far been met with limited success due to the stability of the C-F bonds, and are not suitable for human use. Clinical administration of ion-exchange resins has been shown to be effective but has limited application due to side effects and pharmacological nature of the intervention^22^. We therefore pursued the notable observation of PFAS enrichment within gut bacteria for mechanistic understanding of the bioaccumulation.

Human gut bacteria are known to harbour extensive intra-species genomic and functional variability across individuals, including for bioaccumulation of drugs^23^. To assess the differences in intra- and inter-species bioaccumulation capabilities we tested 89 microbial strains for PFNA sequestration (SI Table 3). These 89 strains covered all major bacterial phyla (Fig. 1b) and included 66 human gut commensal strains, 10 probiotic strains, and 10 bacterial and three yeast strains isolated from kefir. The 66 gut bacterial strains represent, on average, circa 70 % of abundance of the healthy human gut microbiota^10^. PFNA accumulation showed distinct grouping by phylum, with Bacteroidota showing the highest accumulation (Fig. 1b), and a bimodal distribution that could be divided into two groups using a gaussian mixture model, low-accumulating (51 strains), and high-accumulating (38 strains) (SI Fig. 2a; SI Table 6). Low-accumulating strains were predominantly gram-positive (43 out of 51; 84 %) and high-accumulating gram-negative (29 out of 38; 76 %) (SI Fig. 2b). This divide is in contrast with gram-negative bacteria being generally less susceptible to drug and xenobiotic uptake^24,25^. Gram-negative bacteria generally have a higher lipid content (BNID 111942)^26^, which could be a factor underlying PFAS accumulation^27^. However, the divide between gram-negative and gram-positive is not definitive, with notable exceptions such as *E. coli*. Yeasts, which also have a high lipid content^28^ comparable to gram-negative bacteria, also did not show any appreciable PFAS accumulation.

To gain insights into PFAS bioaccumulation, we selected high-accumulating *B. uniformis,* a prevalent gut bacterium^10,11^. PFNA accumulation scaled with cell density, both in non-growing suspension cultures and during bacterial growth (Fig. 1c,d, SI Fig. 2d,e). We next probed kinetics of PFNA uptake using stationary phase cultures or cells suspended in PBS. In both cases bioaccumulation occurred within the timescale of sampling (< 3 min) (Fig. 1e, SI Fig. 2f). This fast kinetics was surprising as previous reports from environmental bacteria isolated from PFAS contaminated sites, *Pseudomonas sp*., showed much lower efficiency with ∼40 % bioaccumulation of perfluorohexane sulfonate (PFHxS) over 5 days, and that following solvent pre-conditioning to facilitate sequestration^29^. Notably, for *B. uniformis*, no PFNA was released back into the supernatant over the course of seven days (Fig. 1e). The fraction of PFAS sequestered was similar, circa 50 % for PFNA, over a broad range of exposed concentrations (0.01-500 μM (4.64 µg/l – 232 mg/l)), both in growing cultures and resting cells (Fig. 1f, SI Fig. 2h,i). The degree of bioaccumulation by *B. uniformis* increased with increasing PFAS chain length, from none for perfluoroheptanoic acid (PFHpA, 7C) to 60 % in the case of perfluorodecanoic acid (PFDA, 10C) (Fig. 1g, SI Fig. 2g). Despite this high degree of bioaccumulation, *B. uniformis* as well as other abundant gut bacterial strains grew well even at high micromolar concentrations (Fig. 1h). Thus, PFAS accumulation reaches a fast equilibrium without inhibiting bacterial growth up to concentrations orders of magnitude higher than known contamination levels.

To investigate how PFAS is bioaccumulated, we first tested whether inactive cell mass (i.e., dead, lysed cells) could bioaccumulate PFAS. In resting cell assays, both live and inactivated *B. uniformis* and *O. splanchnicus* cells bioaccumulated PFOA, PFNA and PFDA to the same extent (∼20 %, ∼55 %, and ∼85 % respectively). In contrast, live *E. coli* bioaccumulated much lower levels of PFOA, PFNA and PFDA (∼5 %, ∼25 % and ∼40 %), while inactivated cells accumulated to a similar degree as *B. uniformis* and *O. splanchnicus* (Fig. 2a). This suggests that PFAS bioaccumulation is not solely a passive phenomenon driven by attachment to membrane lipid bilayers. To test this, we measured bioaccumulation in *E. coli* mutants that lacked one or more genes coding for efflux pump proteins (*imp4213*, Δ*acrA-acrB*, Δ*tolC*) (Fig. 2b). Efflux pumps are a common mechanism used by several bacterial species to reduce intra-cellular concentration of toxic compounds^30,31^. We reasoned that *E. coli* did not bioaccumulate PFNA to the same extent as other tested gram-negative bacteria because it could, at least to some extent, pump out the PFAS. As gene deletions can exhibit different phenotypes in different strain backgrounds^32^, we used two *E. coli* strains, viz. BW25113 and C43 (DE3). Mutants lacking TolC showed a circa 1.5-fold increase in PFDA and circa 5-fold increase in PFNA bioaccumulation (Fig. 2c, SI Fig. 3a). Consistent with the increased bioaccumulation, these mutants also showed increased growth sensitivity (SI Fig. 3b). These results show that *E. coli* strains utilize a TolC dependent mechanism to limit PFAS accumulation.

**Figure 2.**
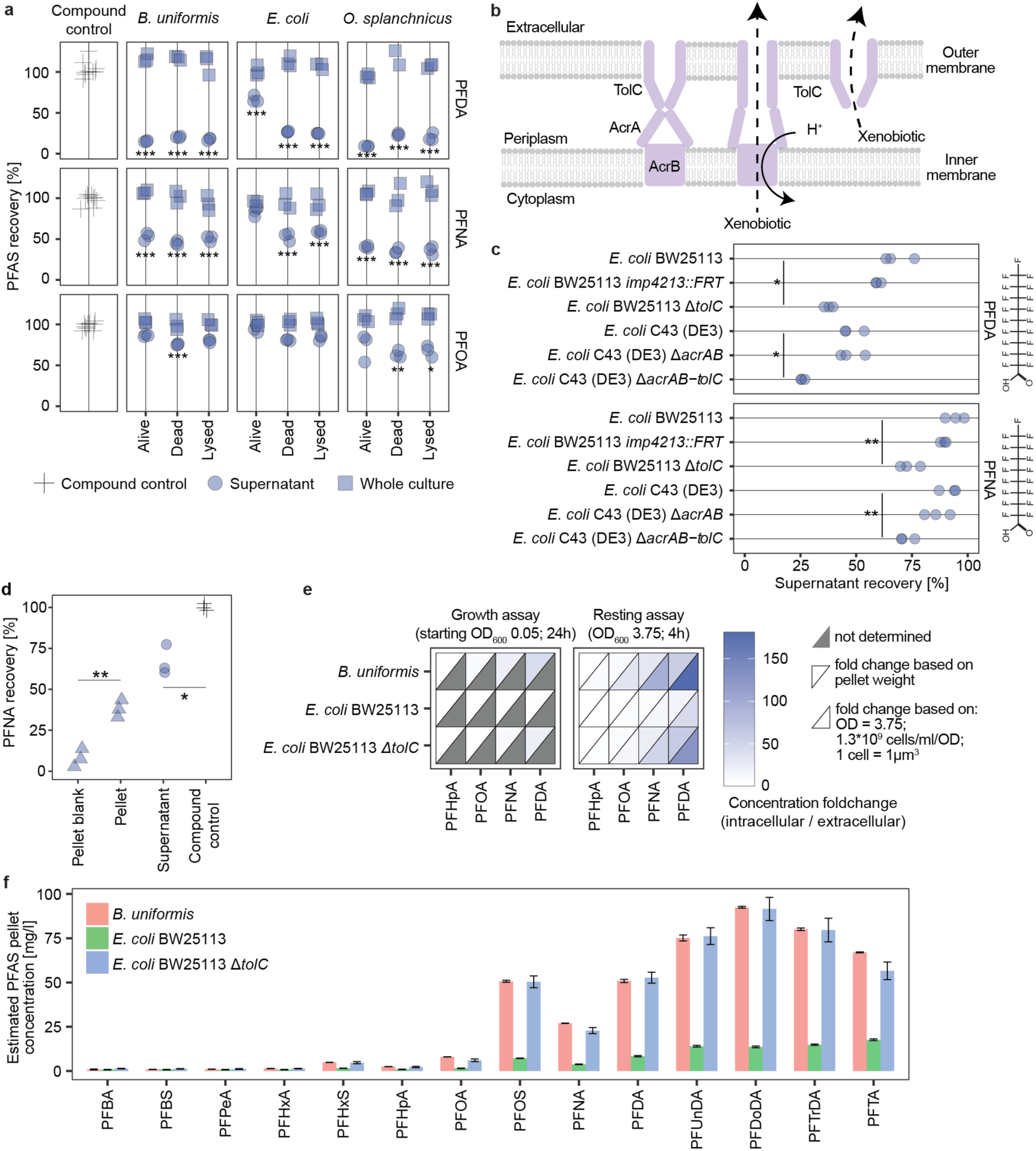
Gut bacteria show a large capacity to concentrate PFAS from their surrounding environment. **a.** FAS bioaccumulation by live, dead (heat-inactivated) and lysed (heat-inactivated + freeze-thawed + sonicated) *B. uniformis*, *E. coli*, and *O. splanchnicus* cultures (OD600 = 3.75) in PBS buffer. * p-value < 0.05; ** p-value < 0.01; *** p-value < 0.001 and > 20 % reduction compared to the compound control; n = 3 technical replicates (SI Table 18). **b.** ArcAB-TolC efflux pump schematic. The resting state of the pump is pictured in the left of the image. When the pump encounters a xenobiotic, it changes conformation to export it out of the cell (middle). TolC can work in combination with a multitude of other pumps and also by itself to pump out xenobiotics from the periplasm (right)^34–36^. **c.** Accumulation of PFDA and PFNA by wild-type *E. coli* strains and corresponding efflux mutants. Efflux mutants *E. coli* BW25113 Δ*tolC* and *E. coli* C43 (DE3) Δ*arcAB-tolC* showed a ∼1.5-fold increase in PFDA and ∼5-fold increase in PFNA bioaccumulation. OD600 = 3.75; exposure concentration = 20 µM (PFNA 9,280 µg/l, PFDA 10,280 µg/l); * p-value < 0.05; ** p-value < 0.01 compared to the corresponding wild-type; n = 3 technical replicates (SI Table 19). **d.** PFNA accumulation by *B. uniformis* at 0.34 nM (160 ng/l) exposure concentration. 37% of PFNA is sequestered from the media into the bacterial pellet corresponding to significant sequestration from the supernatant. * p-value < 0.05; ** p-value < 0.01; n = 3 biological replicates (SI Table 20,21). **e.** Capacity of gut bacteria to concentrate PFAS from the media into the bacterial pellet in a growth assay in mGAM or resting assay in PBS at µM PFAS exposure (PFHpA 1.82 mg/l, PFOA 2.07 mg/l, PFNA 2.32 mg/l, PFDA 2.57 mg/l) (Growth assay: initial OD600 = 0.05, 24 h incubation; resting assay: OD600 = 3.75, 4 h incubation). Concentrating capacity of bacterial strains is an underestimation compared to calculations based on OD600, number of bacterial cells per OD unt (1.3*10^9^ cells/ml/OD) and volume of bacterial cell (1 µm^3^) n = 3 technical replicates (SI Table 23). **f.** PFAS recovery from the bacterial pellets after 1 h exposure in PBS (OD600 = 3.75, PFAS mix of 14 compounds each at a concentration of 1 mg/l). Bars depict the median concentration in the bacterial pellet based on pellet weight determined in b; error bars show standard error; n = 3 technical replicates (SI Table 25).

We next tested whether bacterial cells can bioaccumulate PFAS at very low exposure levels. *B. uniformis* cells were exposed to 0.34 nM (160 ng/l) PFNA, a PFAS concentration observed in water samples across Europe and the US^12,33^ and below the average blood levels^1^. Even at this low exposure level, *B. uniformis* accumulated 37% of PFNA into the bacterial pellet, concordant with depletion from the supernatant (Fig. 2d, SI Fig. 3c). The estimated concentration within the bacterial pellet (wet biomass) was circa 17.7 nM (8200 ng/l), a 50-fold increase (SI Fig. 3d). For 5 µM (PFHpA 1.82 mg/l, PFOA 2.07 mg/l, PFNA 2.32 mg/l, PFDA 2.57 mg/l) exposure, the concentration within the wet pellet of *B. uniformis* and *E. coli* BW25113 Δ*tolC* ranged from circa 13 µM (5 mg/l) for PFHpA, 50 µM (21 mg/l) for PFOA, 130 µM (60 mg/l) for PFNA, to 250 µM (129 mg/l) for PFDA, i.e. 3-, 8-, 25- and 60-fold increase respectively (Fig. 2e, SI Fig. 3e). Since these estimates are based on wet pellet weight, intracellular concentrations will be higher. Considering the estimated number of cells per Optical density (OD) unit (0.6-2 x 10^9^ cells/ml/OD_600_^26^) and 1 µm^3^ cell volume (BNID 101788^26^), the enrichment of PFAS in bacterial cells would be 1.2-7 times higher than in the wet pellet. These results show large capacity of bacterial cells to bioaccumulate and concentrate PFAS from their surrounding environment.

The environmental contamination often contains a mixture of multiple PFAS compounds. We therefore investigated the accumulation capacity of bacteria exposed to a mixture of 14 PFAS (C4-C14, and three sulfonated variants), each at a concentration of 1 mg/l (circa 2µM for PFNA). The results confirm increasing accumulation with increasing chain length (Fig. 2f, SI Fig. 3f). While *E. coli* wild type showed minimal accumulation, *E. coli* Δ*tolC* and *B. uniformis* accumulated almost 100 % of PFAS with chain length above 10C. The percentage of accumulated PFAS for individual compounds is comparable to single compound exposure. Thus, at 1 mg/l per compound, PFAS accumulation seems to be additive across different molecules with minimal mixture effects.

Intra-cellular PFNA concentration in *B. uniformis* at 250 µM (116 mg/l) exposure level would be circa 5-10 mM (2.3-4.6 g/l), well above that of most native metabolites^37^. At these concentrations, the observed degree of bioaccumulation, if considered to be membrane only, would imply one molecule of PFNA per two lipid molecules in the cell membrane, which is physiologically improbable. How do cells cope with such high intra-cellular levels of highly effective surfactants like PFAS and maintain their growth? Towards answering this, we used Transmission Electron Microscopy (TEM) to visualize *B. uniformis, O. splanchnicus, E. coli* wildtype and *E. coli* Δ*tolC* mutant exposed to PFAS. Consistent with their observed levels of bioaccumulation, all cells except *E. coli* wild-type showed disruption and condensation of cytoplasmic content (Fig. 3a,b, SI Fig. 4a-j). Similar morphological features have previously been observed in *Clostridium perfringens* exposed to lauric acid, which also has surfactant properties^38^. We quantified cytoplasmic condensation using automated image processing (Methods, Fig. 3c,d). *B. uniformis* and *O. splanchnicus* showed significantly increased mean pixel intensity and number of condensates upon PFNA/PFDA exposure (Fig. 3e,f, SI Fig. 4k,l). However, TEM imaging can show artefacts due to extensive sample processing. Therefore, to confirm PFNA localisation inside the bacterial cells, we performed cryogenic FIB-SIMS (focused ion beam time-of-flight secondary electron mass spectrometry) imaging of *E. coli ΔtolC* cells exposed to 250 µM PFNA. This pixel-by-pixel three-dimensional analysis of the bacterial cell composition uncovered a strong fluorine signal localised inside the bacterial cells, corresponding to intracellular PFNA accumulation (Fig. 3g-k, SI Fig. 5). In total, 2 biological replicates were analysed, comprising 120 *E. coli ΔtolC* cells exposed to 250 µM PFNA and 44 control cells. All 120 cells exposed to PFNA showed a clear fluorine signal within the bacterial cell, while 43 control cells showed no fluorine signal, and one control cell showed a very low fluorine signal. Further, intracellular PFNA was unevenly distributed with distinct tendency for aggregation. The morphological changes and intracellular aggregation suggest either interaction of PFAS with cytosolic contents, such as proteins and other macromolecules, or phase separation of PFNA and cytosolic contents within the bacterial cell. The containment of the toxicant in these aggregate structures thus appears to be an effective mechanism for the cells to maintain their viability and growth.

**Figure 3.**
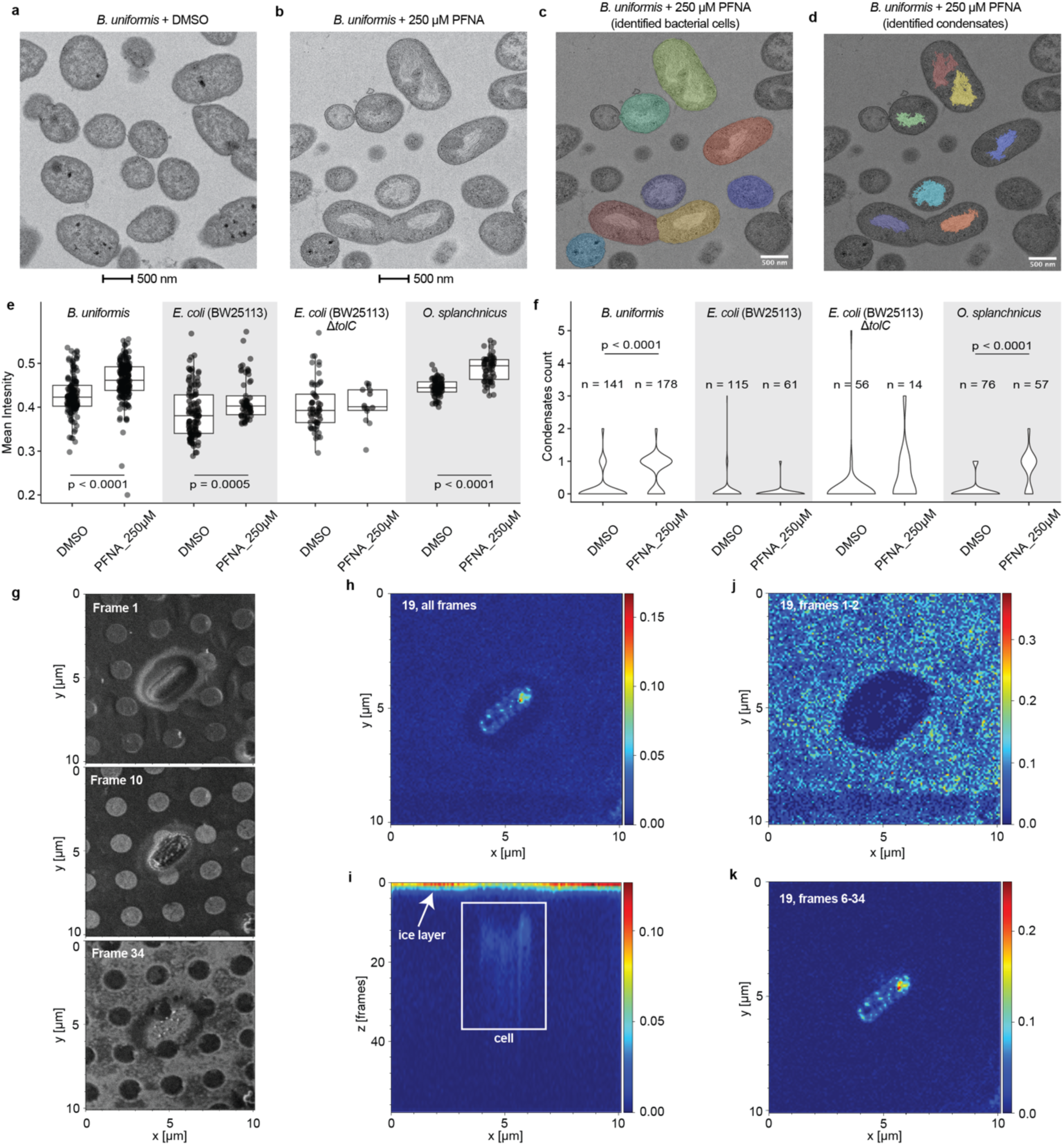
Electron microscopy and imaging mass spectrometry show intra-cellular accumulation of PFAS. **a-f.** Bacteria show distinct morphological features in transmission electron microscopy (TEM). **a,b.** TEM of *B. uniformis* cells grown in mGAM + DMSO (**a**), or 250 μM (116 mg/l) PFNA (**b**) for 24 h. **c,d.** Automatic identification of bacterial cells (**c**) and condensates (**d**) for *B. uniformis* cells grown in mGAM + 250 µM (116 mg/l) PFNA. **e.** Mean pixel intensity of bacterial cells show significant differences between DMSO and PFNA treated cells for *B. uniformis*, *E. coli* BW25113 and *O. splanchnicus* (SI Table 27). **f.** Condensate count per bacterial cell show significant increase in condensates for *B. uniformis* and *O. splanchnicus*, supporting morphological changes through PFNA exposure (SI Table 27). A tomogram for *B. uniformis* cells exposed to 250 µM PFNA confirming that the structures are inside the cells is available as a supplementary video. **g-k.** FIB-SIMS imaging of *E. coli ΔtolC* exposed to 250 µM PFNA. A given area of the sample is imaged by scanning over it repeatedly with a Gallium focused ion beam (FIB) and analysing the chemical composition of the ablated material at each pixel using time-of-flight secondary electron mass spectrometry (ToF-SIMS). As material is continuously removed, the repeated scanning provides three-dimensional information about the sample, hence each 2D image resulting from a full scan of the observed area can be considered as a slice of the sample (frame) within a ‘Z’-stack. This not only allows this method to show the lateral localisation of elements of interest, but also provides depth information. **g.** Secondary electron images (secondary electrons resulting from the FIB scanning) of three different Z-frames of the sample, i.e. at three different positions along the Z-stack, provide spatial images of the imaged cells. While the cells are still fully embedded in ice in the first panel shown (top, frame 1), the second panel (centre, frame 10) shows the interior of the cell and the last panel (bottom, frame 34) features the substrate with the cell almost completely removed. **h-k.** Top (**h, j, k**) views and side view (**i**) of the 3D-stack of SIMS data for mass-to-charge ratio 19, corresponding to fluorine (F^-^). The top views show the lateral distribution (X-Y) of fluorine within the imaged area and confirms its localisation within the cells. The side view (**i**), corresponding to an X-Y-slice through the stack, further corroborates this: for the first few frames, there is fluorine signal from the whole field of view, stemming from the thin ice layer covering the sample. As this is milled away quickly by the Gallium beam, the fluorine signal away from the cells drops to zero within the first frames. As the cells are initially covered in ice, in the first frames, no highly localised fluorine signal is observed from the cells, as demonstrated by the top view generated from the initial frames (**j**). Once the ion beam mills into the cell, fluorine signal from the cell can be seen both in the side view (**i**) and in the top view generated from the corresponding slices (**k**), confirming that the fluorine signal originates from inside the cell.

We next reasoned that if the cells had mechanisms to modulate bioaccumulation and/or to cope with high intra-cellular levels, resistance to inhibitory PFAS exposure would rapidly evolve under natural selection. We therefore evolved, through serial transfer, *B. uniformis*, *B. thetaiomicron*, *P. merdae*, *C. difficile* and *E. coli ΔtolC* in a medium with 500 μM (182 mg/l) PFHpA, 500 μM (207 mg/l) PFOA, 250 μM (116 mg/l) PFNA or 125 μM (64 mg/l) PFDA (SI Fig. 6a). Within 20 transfers (20 days) – corresponding to circa 100 generations – *P. merdae* (PFDA), *B. uniformis* (PFNA, PFDA), and *E. coli ΔtolC* (all tested PFAS) showed between 1.3 and 46 fold increase in growth (SI Fig. 6b-d). Notably, the evolved cells retained the bioaccumulation capability of the respective parental strains (SI Fig. 6e), which is promising for PFAS removal under high and/or chronic exposure environments. Genome sequencing of evolved *B. uniformis* populations identified 55 variants linked to PFNA and PFOA exposure (SI Table 31). More than half of the variants are in non-coding regions, suggesting that changes in gene expression contribute to the adaptation of *B. uniformis* to PFAS.

At the proteomic level, bioaccumulating *E. coli ΔtolC* mutant showed more changes than the non-accumulating wild-type. Most PFNA impacted proteins are either membrane or stress response related (SI Fig. 7a-c). *B. uniformis* also showed changes in membrane related proteins, with the top three being efflux pumps, viz. R9I2M9 efflux transporter RND family, R9I2R1 NodT family efflux transporter, and R9I2L8 hydrophobe/amphiphile efflux-1 (HAE1) family RND transporter (Fig. 4a). Missense variants of the latter RND transporter (R9I2L8) were also identified in all four PFNA evolved *B. uniformis* populations with high (>0.92) allele frequencies. The proteomics and genomic changes together show the involvement of efflux pumps in *B. uniformis* in cellular interaction with PFAS, as we found also for *E. coli*. To identify proteins interacting with PFNA, we used thermal proteome profiling (TPP), which allows proteome wide assessment of structural changes (stabilisation or destabilisation) upon ligand binding^39,40^. We thus compared *E. coli* wild type and *ΔtolC* mutant exposed to PFNA, with the exposure before or after cell lysis, with the latter removing the membrane barrier for PFNA-protein interactions. The bioaccumulating *ΔtolC* mutant featured circa 10-fold more PFNA interacting proteins (SI Table 33). Further, the thermal proteomic responses in the lysate and the live cells are more similar for the mutant than for the wild-type, consistent with increased intracellular PFNA accumulation in the *ΔtolC* mutant (Fig. 4b,c).

**Figure 4.**
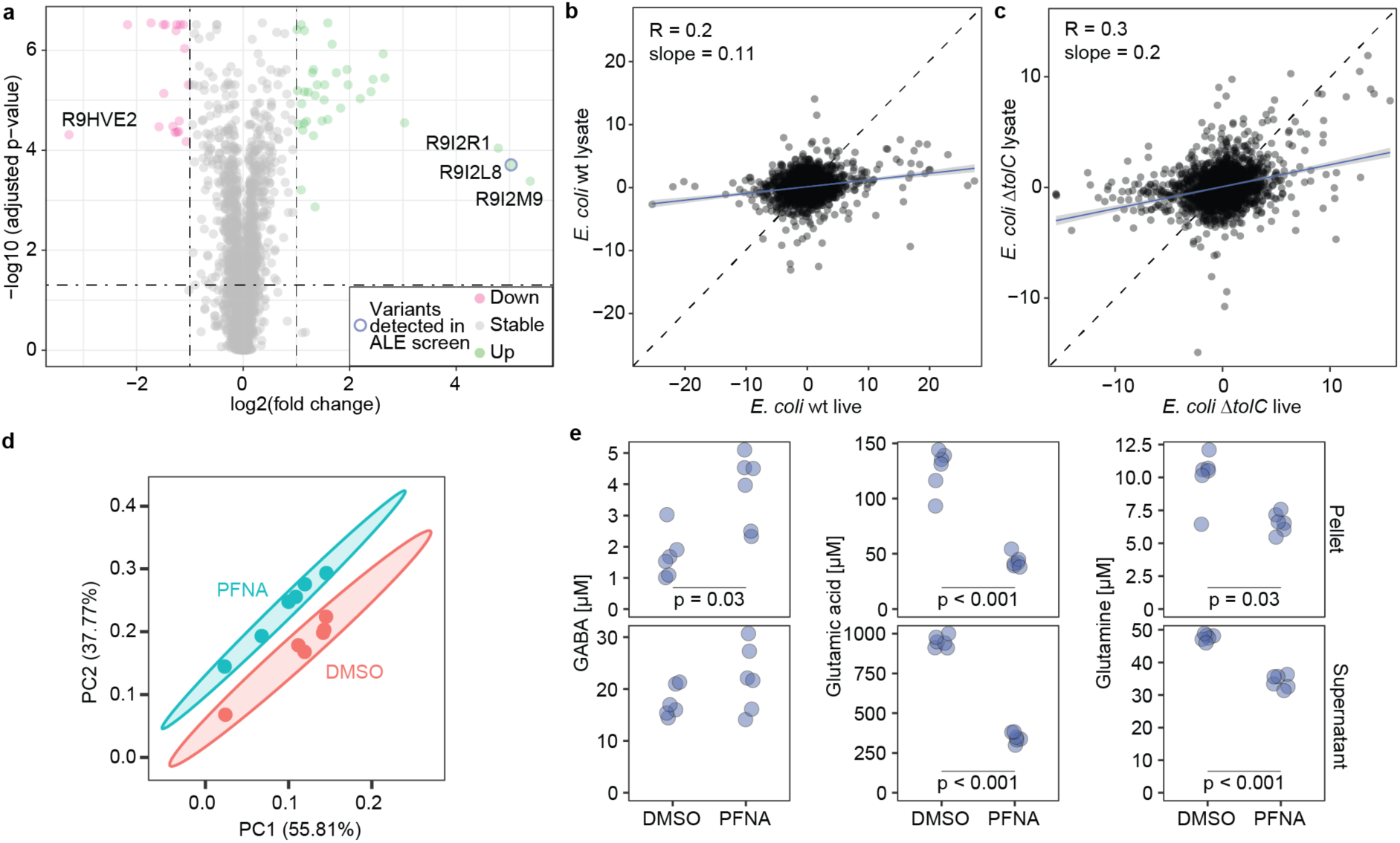
Bioaccumulation of PFNA affects bacterial physiology. **a.** Proteomics analysis show proteins that are differentially abundant between *B. uniformis* treated with 20 µM (9.3 mg/l) PFNA in comparison to DMSO. The red and green dots mark proteins with a log2 abundance ratio >1 or <-1 (i.e., two-fold increase or decrease) and a multiple-testing corrected p-value of less than 0.05; n = 6 biological replicates (SI Table 31,32). **b,c.** TPP analysis of (**b**) *E. coli* BW25113 wild-type (low-PFNA accumulating) and (**c**) *E. coli* BW25113 *ΔtolC* (high-PFNA accumulating). Lysate and live cells incubated with PFNA look more similar for *E. coli* BW25113 *ΔtolC* mutant compared to the wild-type, supporting increased accumulation in *ΔtolC* mutants. Each data point represents the summed log2-fold changes across all temperatures for a specific protein. Black dotted line: diagonal; blue line: linear regression with 95 % confidence interval (SI Table 34). **d.** PCA analysis shows clear distinction between *B. uniformis* pellet samples treated with 20 µM (9.3 mg/l) PFNA compared to the control; n = biological replicates (SI Table 35). **e.** GABA, glutamic acid and glutamine concentration in *B. uniformis* pellet and supernatant samples; n = 6 biological replicates (SI Table 35).

The proteomic changes in efflux pump and other membrane related proteins could be expected to result in altered levels of cellular metabolites. To test this, we used a targeted metabolomics method aimed at broadly conserved metabolites, including amino acids and vitamins. Metabolic changes were observed only for the bioaccumulating strains. *B. uniformis*, one of the highest bioaccumulators, showed distinct metabolic response to PFNA exposure (Fig. 4d,e, SI Fig. 8). Metabolites with altered levels include several amino acids and kynurenine, a metabolite involved in microbiome-host interaction^41,42^. Increased cadaverine and GABA levels, along with decreased glutamate and glutamine levels indicate *B. uniformis* response similar to that observed in acid stress^43–45^.

To determine whether bacterial PFAS bioaccumulation occurs in an *in vivo* context, we tested C57BL/6 mice colonised with a community of 20 human gut bacterial strains (Com20)^46^ (SI Table 37) against germfree controls. The animals were administered a one-time oral dose of PFNA (10 mg/kg body weight) by gavage. Fecal samples were collected over the following 2 days and on day 3 colon and small intestine content samples were taken post-euthanization (Fig. 5a). 17 of the 20 strains colonised the mice and PFNA treatment had no effect on the microbiota composition (SI Fig. 9a,b). Colonized mice showed a significantly higher fecal PFNA excretion at all follow-up time points (3h, p=0.009, fold change, FC=7.9; 1d, p=0.001, FC=2.6; 2d, p<0.0001, FC=2.8; 3d colon, p=0.0004, FC=3.1; 3d small intestine, p=0.007, FC=1.9) (Fig. 5b). Next, we tested whether microbiota constituted of high-versus low-accumulating species would cause difference in fecal PFNA excretion. Mice were either colonized with a community of five high-or five low-PFNA-accumulating strains (LC: *Akkermansia muciniphila, Colinsella aerofaciens, Enterocloster boltae, E. coli, Ruminococcus gnavus*; HC: *Bacteroides fragilis, B. thetaiotaomicron, B. uniformis, Lacrimispora saccharolytica, Phocaeicola vulgatus*, SI Table 37). Similarly to the previous experiment, PFNA treatment had no effect on gut microbiota composition (SI Fig. 9c); however, LC colonised mice showed circa 2-fold higher colonisation compared to HC colonised mice (SI Fig. 9d,e). Per bacterial biomass HC colonised mice showed increased fecal excretion of PFNA on days 1 and 2 (1d, p=0.02, FC=2.4; 2d, p=0.009, FC=3.8) (Fig. 5c). We also carried out the germfree vs. colonised comparison using a 100x lower PFNA dose (0.1 mg/kg BW PFNA), which showed the same trend of increased excretion for the colonized mice (Fig. 5d). Together, the mouse experiments demonstrate the contribution of gut bacterial bioaccumulation to fecal PFAS excretion.

**Figure 5.**
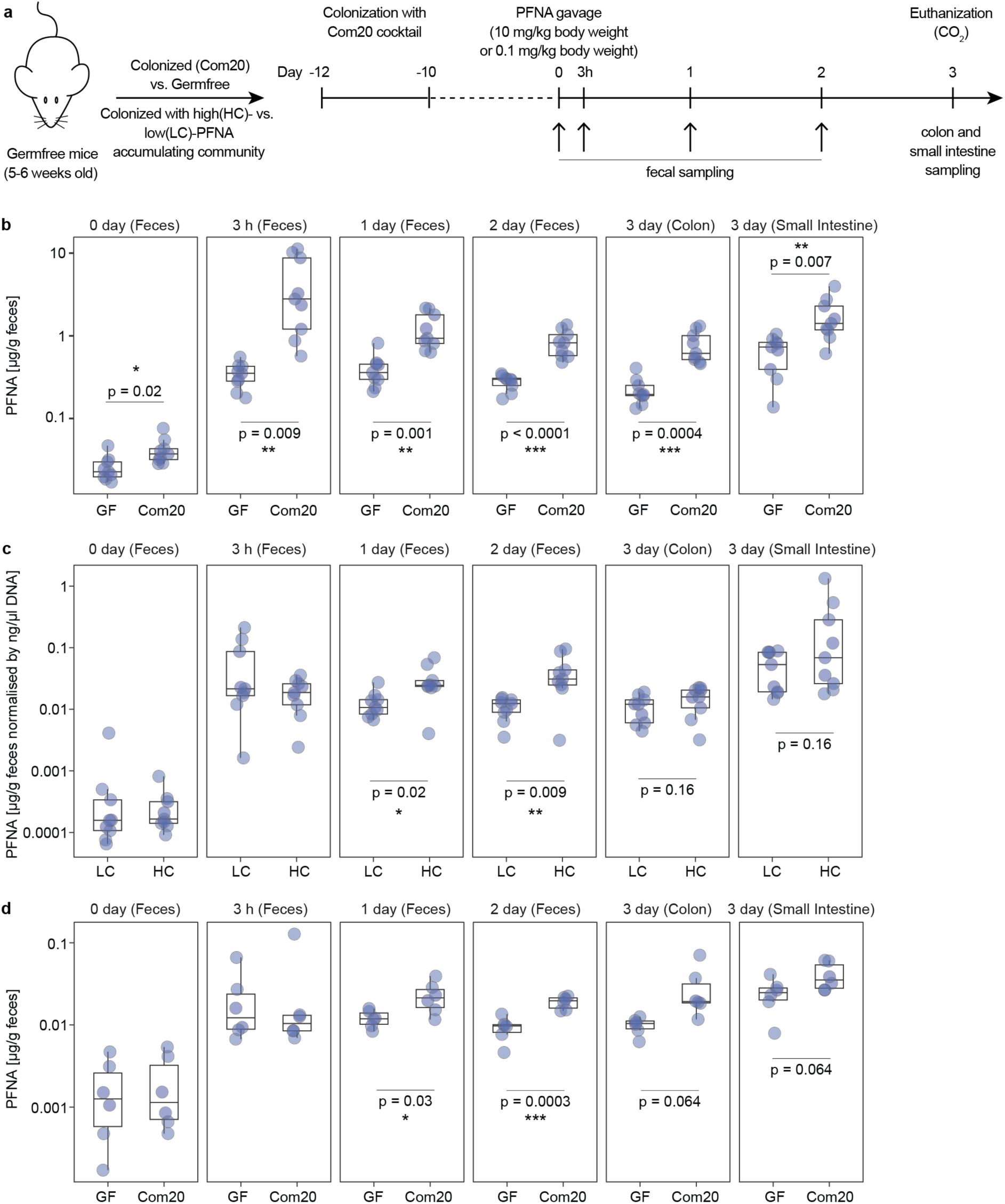
PFNA levels in mouse feces and GI tract are microbiota-dependent. **a.** Experimental setup for comparison of fecal excretion between GF mice and mice colonised with human gut bacteria (Com20) or comparison between mice colonised with high-PFNA accumulating gut bacteria (HC) compared to low-PFN accumulating gut bacteria (LC). **b.** Mice colonised with a community of 20 human gut bacterial strains (Com20) show higher PFNA excretion after 10 mg/kg body weight PFNA exposure compared to germfree (GF) controls (SI Table 38). **c.** Mice colonised with a community of high-PFNA accumulating gut bacteria (HC) show higher PFNA excretion after 10 mg/kg body weight PFNA exposure compared to mice colonised with low-PFNA accumulating gut bacteria (LC) (SI Table 39). All y-axes are on log10 scale. **d.** Mice colonised with a community of 20 human gut bacterial strains (Com20) show higher PFNA excretion after 0.1 mg/kg body weight PFNA exposure compared to germfree (GF) controls (SI Table 40).

Overall, our study discovers the remarkable capacity of human gut bacteria to accumulate PFAS. The internalisation of PFAS is supported both by analytical and biological evidence (multi-omics analyses, imaging, phylogenetic grouping of accumulators, and difference between wild-type and efflux-pump mutant). There is no correlation between PFAS accumulation by gut bacteria and the previously observed drug bioaccumulation ^23^, suggesting independence of the two phenomena. The PFAS bioaccumulation is notable because of the high capacity, rapid kinetics, and fundamental implications for understanding how PFAS interacts with biological systems. While we have identified efflux-pump based export mechanism, additional exporters may be involved, and it remains unclear how PFAS enters the cell. Intracellular PFAS distribution as determined by FIB-SIMS shows distinct aggregation inside the bacterial cells. However, its (re-)distribution following cell division remains an open question. FIB-SIMS analysis of growing cells in presence of PFAS could help elucidate re-distribution of PFAS during cell division. In addition, this technology could help determine differences in intracellular PFAS localisation for different PFAS compounds.

The data from our gnotobiotic mouse studies shows that gut bacteria accumulate PFAS also in an in vivo context. Yet, these experiments were done using a one-time high dosage while most populations experience low but chronic exposure. Therefore, cohort studies tracking PFAS intake, body levels, excretion, and microbiota composition over a prolonged period will be needed to understand PFAS toxicokinetics. Our results bring forward the interaction of PFAS with gut bacteria opening the possibilities for the development of biological approaches for PFAS removal.

## Acknowledgements

We thank JL Gonzalez for helpful discussions on TEM imaging, E Kafkia for advice on mass-spectrometry, and N Typas and B Luisi labs for providing the *E. coli* efflux pump mutants. A.E.L., L.P.B and A.K.R. acknowledge funding by the National Institute for Health and Care Research (NIHR) Health Protection Research Units in Chemical and Radiation Threats and Hazards and/or Environmental Exposures and Health. K.R.P. and A.E.L. acknowledge funding by UK Medical Research Council (project no. MC_UU_00025/11).

## Author contributions

KRP and AEL conceived and planned the overall study. AEL performed the microbiological and mass-spectrometry experiments and data analysis. SK contributed to mass-spectrometry experiments, mouse fecal sample processing and data analysis. RG contributed to the laboratory evolution experiment and metabolomics analysis and performed and analysed the genome sequencing for evolved populations. SB advised on and contributed to microbiological experiments. IR advised on *E. coli* mutant experiments. AKR and LPB performed an independent LCMS analysis of the 160 ng/l PFNA experiment samples. MGM and NM performed the electron microscopy experiments and image acquisition. LV and AZ contributed to TEM image analysis. BR performed and analysed the proteomics experiment. DP and MS performed and analysed the TPP experiments. AG, LMichaelis and LMaier planned and performed the animal experiments. NB, JYP and JEDT advised on design of animal experiments and contributed to animal sample processing. HOzgur and VB performed 16S sequencing. HOchner and TAMB carried out the cryogenic FIB-SIMS analysis. AEL and KRP wrote the manuscript. All authors commented on the manuscript.

## Methods

### Bacterial strains and cultivation

Strains were selected to represent prevalent and abundant members of the healthy human gut microbiota^10,11^ (SI Table 1). *E. coli* mutants were obtained from Typas lab (EMBL Heidelberg, BW25113 wild-type, BW25113 Δ*tolC,* BW25113 *imp4213::FRT*) and Luisi lab (University of Cambridge, C43 (DE3) wild-type, C43 (DE3) Δ*acrAB-tolC,* C43 (DE3) Δ*acrAB*). All bacterial experiments were performed in an anaerobic chamber (Coy Laboratory Products) filled with 2 % hydrogen and 12 % carbon dioxide in nitrogen. The chamber was equipped with a palladium catalyst system for oxygen removal, a dehumidifier, and a hydrogen sulfide removal system. Bacteria were grown at 37 °C in modified Gifu anaerobic medium (mGAM, HyServe, Germany, produced by Nissui Pharmaceuticals), prepared according to the instructions from the manufacturer and sterilised by autoclaving. Bacteria for starting cultures were grown for one or two days (depending on growth rate) in 10 ml of media in 15 ml plastic tubes, which were inoculated directly from frozen glycerol stocks. Cultures were then diluted 100-fold and incubated again for the same amount of time before starting the experiments. Unless otherwise specified, the screening plates/tubes with cultivation medium were prepared the day prior at 2x compound concentration (2 % DMSO) and placed into the chamber over night to ensure anaerobic conditions for inoculation. Inoculation was performed 1:1 with a bacterial culture and plates were sealed with AlumaSeal II film (A2350-100EA) to avoid evaporation during incubation.

### Community-based screening approach (SI Fig. 1a,b)

On the day of the screen communities were assembled by pooling together second passages of individual strains according to their OD_600_ values. Assembled communities were then centrifuged at 25 °C and 4000 rpm for 15 min and the pellet was resuspended in PBS buffer to create a community with OD_600_ of 7.5. Each well was inoculated 1:1 with community in PBS to reach a starting OD_600_ of 3.75 and a compound concentration of 20 µM (1 % DMSO). For compound control wells bacteria-free PBS was added to the respective wells. Plates were incubated at 37 °C for 4 h, after which they were centrifuged at 21 °C and 4000 rpm for 15 min. The supernatant and compound controls were transferred to fresh 96-well plates and stored at −80 °C until extraction.

### Single strain-xenobiotic screen (Fig. 1a)

Ten compounds selected based on the ‘Community-based screening approach’ were tested for sequestration by individual strains. On the day of the screen each well was inoculated 1:1 with a second passage culture to reach a starting OD_600_ of 0.05 and a compound concentration of 20 µM (1 % DMSO). For compound control wells bacteria-free mGAM was added to the respective wells. Plates were incubated at 37 °C for 24 h, after which they were removed from the anaerobic chamber for sample collection. Whole culture, supernatant and compound control samples were collected and stored at −80 °C until extraction.

### PFAS bioaccumulation analysis

#### *Resting cell assay* (Fig. 1b,c,g; Fig. 2c; SI Fig. 2a-d,g,I; SI Fig. 3a)

Each well was inoculated 1:1 with a culture in PBS to reach the desired starting OD_600_ of 3.75, unless otherwise specified. For compound control wells bacteria-free PBS was added to the respective wells. Samples were incubated at 37 °C for 4 h, after which they were removed from the anaerobic chamber for sample collection. Whole culture, supernatant and compound control samples were collected and stored at −80 °C until extraction.

#### Growth assay (**Fig. 1f**; SI Fig. 2h)

On the day of the screen each well was inoculated 1:1 with a second passage culture to reach a starting OD_600_ of 0.05. For compound control wells bacteria-free mGAM was added to the respective wells. Samples were incubated at 37 °C for 24 h, after which they were removed from the anaerobic chamber for sample collection. Whole culture, supernatant and compound control samples were collected and stored at −80 °C until extraction.

#### PFAS time-course experiment with growing B. uniformis culture (Fig. 1d; SI Fig. 2e)

On the day of the screen each tube was inoculated 1:1 with a second passage culture to reach a starting OD_600_ of 0.05 and a concentration of 20 µM (9.3 mg/l) PFNA. Samples were incubated at 37 °C for 11 h. Bacteria free mGAM was added to compound control samples. OD_600_ was measured every hour and supernatant, whole culture and compound control samples for PFNA analysis were collected also every hour and stored at −80 °C until extraction.

#### PFAS time-course experiment with resting B. uniformis culture (**Fig. 1e**)

On the day of the screen each well was inoculated 1:1 with a culture in PBS to reach a starting OD_600_ of 3.75 and a concentration of 20 µM (9.3 mg/l) PFNA. Samples were incubated at 37 °C. Bacteria free PBS was added to compound control samples. Supernatant, pellet, whole culture and compound control samples for PFNA analysis were collected at 0h, 4h, 1d, 2d, 4d, and 7d and stored at −80 °C until extraction.

#### PFAS time-course experiment with stationary phase B. uniformis culture (SI Fig. 2f)

1.5 ml of stationary second passage cultures of *B. unifomis* or pure mGAM were spiked with 15 μl of 2 mM PFNA in DMSO (final concentration 20 µM (9.3 mg/l) PFNA, 1 % DMSO). Whole culture, supernatant and compound control samples were collected at 0, 15, 30 and 60 min and stored at −80 °C until extraction.

#### PFAS accumulation in live, heat-inactivated, and lysed bacterial cultures (Fig. 2a)

On the day of the screen each well was inoculated 1:1 with an alive, heat-inactivated or lysed culture in PBS to reach a starting OD_600_ of 3.75. Second passage cultures were spun down, and the pellet was resuspended in PBS to an OD_600_ of 7.5. Each culture was split up into 3 aliquots: alive, heat-inactivated, lysed cultures. Live cultures were used as is. Bacteria were heat-inactivated at 70 °C for 40 min and lysed cultures were additionally freeze-thawed three times and sonicated for 3 min. After adding the respective cultures or bacteria-free PBS to the respective wells the plates were sealed and incubated at 37 °C. After 4 h whole culture, supernatant and compound control samples were collected and stored at −80 °C until extraction.

#### Low concentration experiment (160 ng/l) (Fig. 2d; SI Fig. 3c,d)

Three biological replicates of *B. unifprmis* were grown over the course of two days. On the day of the experiment the OD_600_ of the second passage cultures was measured, and the equivalent of 2x 0.5 l of OD_600_ of 3.75 of each replicate was centrifuged 4000 rpm for 15 min and the supernatant removed. One pellet per biological replicate was kept as negative control and stored at −80 °C until extraction. The other pellet was resuspended in 0.5 l of 160 ng/l PFNA in PBS and incubated at 37 °C for 1 h in HDPE bottles (Buerkle 10531712). 3x 0.5 l of 160 ng/l PFNA in PBS without bacterial pellet was used as the compound control and incubated for the same amount of time. For collection samples were removed from the anaerobic chamber and centrifuged at 4000 rpm for 15 min. Supernatant and pellet samples were stored separately at −80 °C until extraction.

#### PFHpA, PFOA, PFNA and PFDA pellet recovery (Fig. 2e, SI Fig. 3e)

To determine pellet concentration based on pellet weight across four PFAS compounds (PFHpA, PFOA, PFNA, PFDA) a resting cell assay and growth assay was conducted at an exposure concentration of 5 µM in 0.5 ml volume. On the day of the screen each well was inoculated 1:1 with a second passage culture to reach a starting OD_600_ of 0.05 in mGAM or a starting OD_600_ of 3.75 in PBS and a starting concentration of 5 µM of the respective PFAS compound (0.5 ml total volume). Samples were incubated at 37 °C for 24 h (mGAM) or 37 °C for 4 h (PBS), after which they were removed from the anaerobic chamber for sample collection. Supernatant was collected for LCMS analysis, pellets were weighed and also collected for LCMS analysis. In the resting assay *B. uniformis* and *E. coli* pellets weighed ∼8 and ∼5 mg respectively, meaning bacterial cells only contribute circa 1-2 % to total culture weight/volume.

#### Accumulation of a mix of 14 PFAS compounds (Fig. 2f; SI Fig. 3f)

On the day of the screen each well was inoculated 1:1 with a culture in PBS to reach a starting OD_600_ of 3.75 (1 mg/l for each PFAS compound: PFBA, PFBS, PFPeA, PFHxA, PFHxS, PFHpA, PFOA, PFOS, PFNA, PFDA, PFUnDA, PFDoDA, PFTrDA, PFTA). For compound control wells bacteria-free PBS was added to the respective wells. Samples were incubated at 37 °C for 4 h, after which they were removed from the anaerobic chamber for sample collection. Whole culture, supernatant, pellet and compound control samples were collected and stored at −80 °C until extraction.

#### PFNA and PFDA accumulation by evolved populations (SI Fig. 5e)

20 µl of a second passage culture was added to 380 µl PFAS in mGAM to reach a starting concentration of 20 µM PFNA or PFDA. Bacteria free mGAM was added to compound control samples. Samples were incubated at 37 °C for 24 h, after which whole culture, compound control and supernatant samples were collected for LCMS analysis.

### Standard compounds

For all measured compounds, including PFHpA, PFOA, PFNA and PFDA, pure standards were obtained from Sigma Aldrich (Merck KGaA, Darmstadt, Germany). All compounds were dissolved in DMSO, with the exception of NMOR, MDMA and methamphetamine, which came dissolved in methanol, and cocaine and heroin, which came dissolved in acetonitrile. A mix containing 14 PFAS compounds was purchased from Agilent (ITA-70). Labelled standards of PFNA and a mix of 13 PFAS standards were purchased from Cambridge Isotope Laboratories and Greyhound Chromatography (CLM-8060-1.2, MPFAC-C-ES).

### PFAS solubility

For all PFAS assays, either mGAM or PBS with 1% DMSO was used. To test whether solubility of PFAS compounds could affect our assays, we measured solubility of PFNA and PFDA in mGAM (1% DMSO), PBS (1% DMSO) and 80% methanol (1% DMSO). PFNA was soluble in all conditions up to 500 µM, while PFDA was soluble up to 500 µM in 80% methanol (1% DMSO) and mGAM (1% DMSO), and up to 100 µM in PBS (1% DMSO) (SI Fig. 10a).

### Sample extraction for LC-MS/MS

#### Bacterial samples

For the ‘Community-based screening approach’ 70 μl supernatant was extracted with 140 μl of ice-cold methanol:acetonitrile (1:1) containing the internal standard (20 µM caffeine, 60 µM ibuprofen) and incubated at 4 °C for 30 min. For the ‘Single strain-xenobiotic screen’ 50 μl sample was extracted with 200 μl ice-cold methanol:acetonitrile (1:1) containing internal standard (100 µM caffeine, 60 µM ibuprofen) and incubated at 4 °C for 15 min. For other follow-up *in vitro* assays 50 μl sample was extracted with 200 μl ice-cold methanol:acetonitrile (1:1) containing internal standard (60 µM ibuprofen or 20 µM caffeine) and incubated at 4 °C for 15 min. For the PFAS-mix assay all samples were diluted 1:1 with water and 10 µl diluted sample was extracted with 90 µl ice-cold methanol:acetonitrile (1:1) containing internal standard (final IS concentration 20 µg/l for each internal standard) 4 °C for 15 min. Sample plates were then centrifuged at 4 °C and 4000 rpm for 10 min. Supernatants were transferred to 96-well plates for LC-MS analysis. Samples for concentration calibration and bacteria-free compound controls were processed in the same way.

#### Sample extraction per EPA Draft Method 1633 (Fig. 2d; SI Fig. 3c,d)

Supernatant, and pellet samples were extracted according to the EPA 3^rd^ Draft Method 1633 for analysis of PFAS in aqueous, solid, biosolid and tissue samples^47^ and the corresponding Agilent application note^48^. In short, 500 ml supernatant or compound control samples were weighed and spiked with a known concentration of an internal standard (1 ml of 50 µg/l 13C9-PFNA). Pellet samples underwent three freeze-thaw cycles to ensure bacterial lysis. Samples were weighed, resuspended in 20 ml of 0.3 % methanolic ammonium hydroxide, spiked with a known concentration of an internal standard (1 ml of 50 µg/l 13C9-PFNA) and shaken for 30 min. Samples were then centrifuged at 2800 rpm for 10 min and the supernatant was transferred to a fresh sample tube. The remaining pellet was resuspended in 15 ml of 0.3 % methanolic ammonium hydroxide, shaken for 30 min and centrifuged again. The supernatant was added to the same collection tube and the pellet underwent the same process again using 10 ml of 0.3 % methanolic ammonium hydroxide. To the combined supernatants from each pellet circa 10 mg carbon (Agilent 5982-4482) was added. Samples were hand shaken for up to 5 min and then centrifuged for 10 min at 2800 rpm. The supernatants were collected in a fresh sample tube. Agilent solid-phase extraction (SPE) cartridges (Agilent 5610-2150) were loaded with silanized glass wool (Agilent 8500-1572) fitted with large volume adapters (Agilent 12131012 and 12131001) and conditioned with 15 ml 1% methanolic ammonium hydroxide followed by 5 ml of 0.3 M formic acid. Samples were then loaded into SPE cartridges and set to a low flow rate of circa 5 ml/min, followed by a rinse with 10 ml reagent water and 5 ml of 1:1 0.1 M aqueous formic acid:methanol, before being dried under the vacuum. Sample bottles were rinsed and eluted with 5 ml of 1% methanolic ammonium hydroxide. 25 µl acetic acid was added to each sample and vortexed. To each supernatant and compound control eluate 10 mg carbon (Agilent 5982-4482) was added, samples were handshaken for up to 5 min and then centrifuged for 10 min at 2800 rpm. All samples were filtered through a nylon syringe filter (Agilent 9301-6476, 5190-5092) into a collection tube for LCMS analysis.

#### Mouse fecal samples

Frozen fecal samples were weighed out into tubes with beads and 250 μl extraction buffer (methanol + 0.05 KOH + 15 μM caffeine (Fig. 6b) or 0.01 mg/l M9PFNA (Fig. 6c,d)) was added. Tubes were then homogenised at 1500 rpm for 10 min followed by centrifugation at 14,000 rpm and 4 °C for 5 min. 20 μl supernatant was added to 80 μl water + 0.1 % formic acid, vortexed, incubated at 4 °C for 15 min and centrifuged at 14,000 rpm and 4 °C for 5 min. The supernatant was transferred to LCMS vials with inserts. Samples for concentration calibration were processed in the same way.

### LC-MS/MS (QTOF) xenobiotic measurements

#### QTOF parameters

In short, LC-MS analysis was performed on an Agilent 1290 Infinity II LC system coupled with an Agilent 6546 LC/Q-TOF (Agilent). The QTOF MS scan was operated in positive or negative MS mode using four different collision energies (0V, 10V, 20V, 40V) (30-1500 m/z), depending on the xenobiotic targeted for measurement (SI Table 2). The source parameters were as follows: gas temperature: 200 °C, drying gas: 9 L/min, nebulizer: 20 psi, sheath gas temperature: 400 °C, sheath gas flow: 12 L/min, VCap: 3000 V, nozzle voltage: 0 V, fragmentor: 110 V, skimmer 45 V, Oct RF Vpp: 750 V. The online mass calibration was performed using a reference solution (positive: 121.05 and 922.01 m/z; negative: 112.99 and 1033.99 m/z).The compounds were identified based on their retention time, accurate mass and fragmentation patterns. For all measured compounds pure standards were used for method development, compound identification and calibration. Five different LC-methods were applied.

#### 15 min reverse-phase LC-method used with QTOF in positive ionization mode (’Community-based screening approach’ SI Fig. 1)

The separation was performed using a ZORBAX RRHD Eclipse Plus column (C18, 2.1 × 100 mm, 1.8 μm; Agilent 858700-902) with a ZOBRAX Eclipse Plus (C18, 2.1 × 5 mm, 1.8 μm; Agilent 821725-901) guard column at 40 °C. The multisampler was kept at a temperature of 4 °C. The injection volume was 1 μL and the flow rate was 0.4 mL/min. The mobile phases consisted of A: water + 0.1 % formic acid + 5 mM ammonium formate; B: methanol + 0.1 % formic acid + 5 mM ammonium formate. The 15 min gradient started with 5 % solvent B, which was increased to 30 % by 1 min and then further increased to 100 % by 7 min and held for 3 min, before returning to 5 % solvent B for a 5 min re-equilibration.

#### 10 min reverse-phase dual-pump LC-method used with QTOF in positive ionization mode (Fig. 1a,b)

The separation was performed using two ZORBAX RRHD Eclipse Plus column (C18, 2.1 × 100 mm, 1.8 μm; Agilent 858700-902) with the ZOBRAX Eclipse Plus (C18, 2.1 × 5 mm, 1.8 μm; Agilent 821725-901) guard columns at 40 °C. The multisampler was kept at a temperature of 4 °C. The injection volume was 1 μL and the flow rate was 0.4 mL/min. The mobile phases consisted of A: water + 0.1 % formic acid + 5 mM ammonium formate; B: methanol + 0.1 % formic acid + 5 mM ammonium formate. The 10 min gradient started with 5 % solvent B, which was increased to 30 % by 1 min and then further increased to 100 % by 7 min and held for 1.7 min, before returning to 5 % solvent B at 8.8 min, which was held until 10 min. The re-equilibration gradient started with 5 % solvent B, which was then ramped up to 100 % solvent B at 0.1 min and held until 4 min before returning to the starting condition of 5 % solvent B at 4.1 min.

#### 13 min reverse-phase LC-method used with QTOF in negative ionization mode (’Community-based screening approach’ SI Fig. 1)

The separation was performed using a ZORBAX RRHD Eclipse Plus column (C18, 2.1 × 100 mm, 1.8 μm; Agilent 858700-902) with a ZOBRAX Eclipse Plus (C18, 2.1 × 5 mm, 1.8 μm; Agilent 821725-901) guard columns at 40 °C. The multisampler was kept at a temperature of 4 °C. The injection volume was 1 μL and the flow rate was 0.4 mL/min. The 13 min gradient started with 35 % solvent B, which was increased to 100 % by 9 min and held for 1 min, before returning to 35 % solvent B for a 3 min re-equilibration. Mobile phases for BPA, BPB, BPF and catechol analysis consisted of A: water and B: methanol; Mobile phases for BPAF, BPS, PFNA, PFOA, patulin and 2-nitrofluorene analysis consisted of A: water + 5 mM ammonium acetate + 0.03 % acetic acid and B: methanol + 5 mM ammonium acetate + 0.03 % acetic acid.

#### 10 min reverse-phase dual-pump LC-method used with QTOF in negative ionization mode (Fig. 1a,b,e,g; Fig. 2a,c; SI Fig. 2g,i; SI Fig. 3a)

The separation was performed using two ZORBAX RRHD Eclipse Plus column (C18, 2.1 × 100 mm, 1.8 μm; Agilent 858700-902) with the ZOBRAX Eclipse Plus (C18, 2.1 × 5 mm, 1.8 μm; Agilent 821725-901) guard columns at 40 °C. The multisampler was kept at a temperature of 4 °C. The injection volume was 1 μL and the flow rate was 0.4 mL/min. The mobile phases consisted of A: water + 5 mM ammonium acetate + 0.03 % acetic acid; B: methanol + 5 mM ammonium acetate + 0.03 % acetic acid. The 10 min gradient started with 35 % solvent B, which was increased to 100 % by 7 min and held for 1.7 min, before returning to 35 % solvent B at 8.8 min, which was held until 10 min. The re-equilibration gradient started with 35 % solvent B, which was then ramped up to 95 % solvent B at 0.1 min and held until 4 min before returning to the starting condition of 35 % solvent B at 4.1 min.

#### 2 min reverse-phase LC-method used with QTOF in negative ionization mode (SI Fig. 5e)

The separation was performed using a ZORBAX RRHD Eclipse Plus column (C18, 3.0 × 50 mm, 1.8 μm; Agilent 959757-302) at 40 °C. The multisampler was kept at a temperature of 4 °C. The injection volume was 1 μL and the flow rate was 0.8 mL/min. The mobile phases consisted of A: water + 5 mM ammonium acetate + 0.03 % acetic acid; B: methanol + 5 mM ammonium acetate + 0.03 % acetic acid. The 2 min gradient started with 30 % solvent B, which was increased to 100 % by 0.5 min and held until 1 min, before returning to 30 % solvent B at 1.1 min until 2 min.

### LC-MS/MS (QQQ) PFAS measurements

#### QQQ parameters

In short, LC-MS/MS analysis was performed on an Agilent 1290 Infinity II LC system coupled with an Agilent 6470 triple quadrupole (Agilent). The QQQ was operated in Dynamic MRM mode. The source parameters were as follows: gas temperature: 300 °C, gas flow: 10 L/min, nebulizer: 50 psi, sheath gas temperature: 300 °C, sheath gas flow: 11 L/min, VCap: 3500 V (positive) or 3000 V (negative), nozzle voltage: 2000 V (positive) or 500 V (negative). The transitions for PFAS can be found in Table 1. Pure standards were used for method development, compound identification and calibration.

**Table 1.**
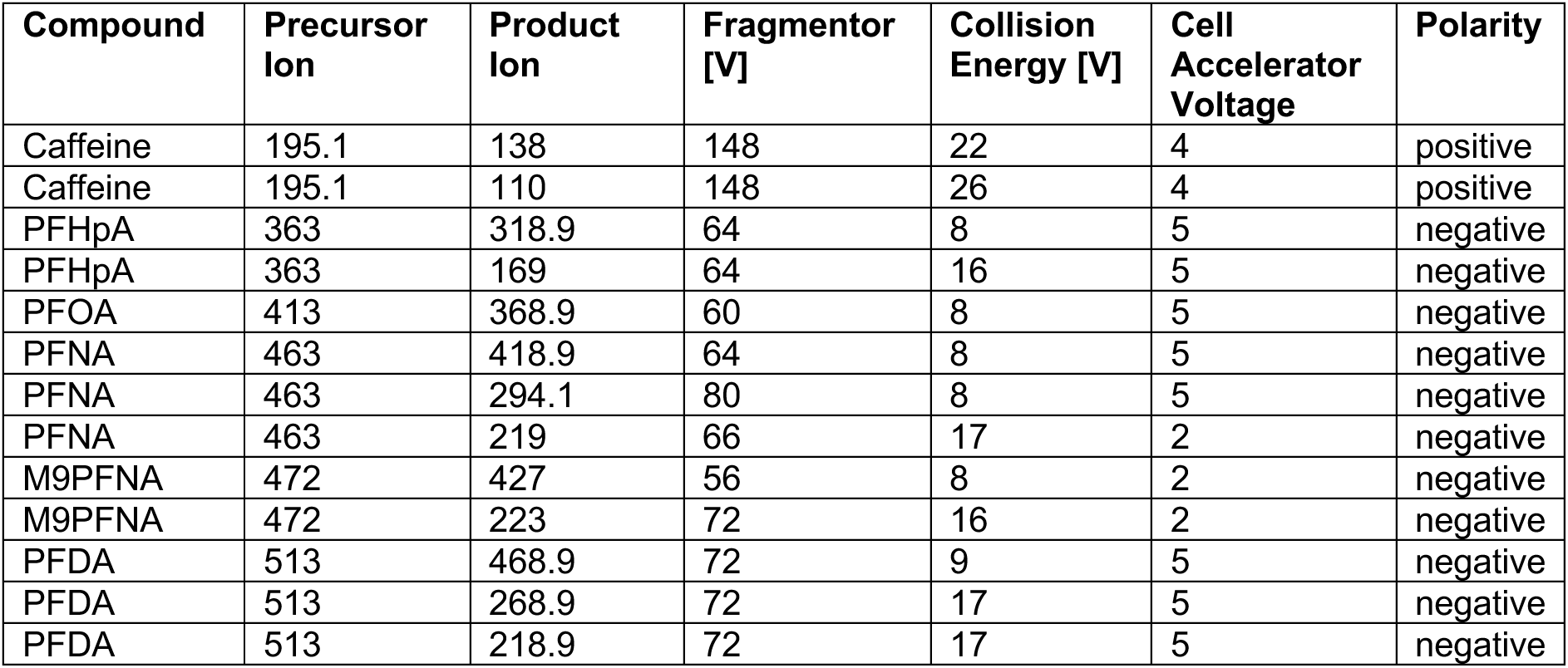
dMRM transitions monitored.

| Compound | Precursor Ion | Product Ion | Fragmentor [V] | Collision Energy [V] | Cell Accelerator Voltage | Polarity |
| --- | --- | --- | --- | --- | --- | --- |
| Caffeine | 195.1 | 138 | 148 | 22 | 4 | positive |
| Caffeine | 195.1 | 110 | 148 | 26 | 4 | positive |
| PFHpA | 363 | 318.9 | 64 | 8 | 5 | negative |
| PFHpA | 363 | 169 | 64 | 16 | 5 | negative |
| PFOA | 413 | 368.9 | 60 | 8 | 5 | negative |
| PFNA | 463 | 418.9 | 64 | 8 | 5 | negative |
| PFNA | 463 | 294.1 | 80 | 8 | 5 | negative |
| PFNA | 463 | 219 | 66 | 17 | 2 | negative |
| M9PFNA | 472 | 427 | 56 | 8 | 2 | negative |
| M9PFNA | 472 | 223 | 72 | 16 | 2 | negative |
| PFDA | 513 | 468.9 | 72 | 9 | 5 | negative |
| PFDA | 513 | 268.9 | 72 | 17 | 5 | negative |
| PFDA | 513 | 218.9 | 72 | 17 | 5 | negative |

*2 min reverse-phase LC-method used with QQQ (Fig. 1b,c,d,f; Fig. 3b; SI Fig. 3e)*

#### 2 min reverse-phase LC-method used with QQQ (Fig. 1b,c,d,f; Fig. 3b; SI Fig. 3e)

The separation was performed using a ZORBAX RRHD Eclipse Plus (C18, 3 × 50 mm, 1.8 μm; Agilent Agilent 959757-302) or a Poroshell 120 EC-C18, 1.9um, 2.1 x 50mm (Agilent 699675-902) at 40 °C. The multisampler was kept at a temperature of 4 °C. The injection volume was 1 μL and the flow rate was 0.8 ml/min. The mobile phases consisted of either A: water + 5mM ammonium acetate + 0.03% acetic acid and B: methanol + 5mM ammonium acetate + 0.03% acetic acid, or A: water + 0.1 % formic acid and B: acetonitrile + 0.1 % formic acid. The 2 min gradient started with 30 % solvent B, which was increased to 100 % by 0.5 min and held until 1 min, before returning to 30 % solvent B at 1.05 min and held until 2 min.

#### 10 min reverse-phase LC-method used with QQQ for mouse fecal sample analysis (Fig. 5)

The separation was performed using a ZORBAX RRHD Eclipse Plus column (C18, 2.1 × 100 mm, 1.8 μm; Agilent 858700-902) with a ZOBRAX Eclipse Plus (C18, 2.1 × 5 mm, 1.8 μm; Agilent 821725-901) guard column at 40 °C. The multisampler was kept at a temperature of 4 °C. The injection volume was 2-10 μl (Fig. 6b: 2 µl; Fig. 6c: 10 µl; Fig. 6d: 5 µl) and the flow rate was 0.4 mL/min. The mobile phases consisted of A: water + 0.1 % formic acid; B: acetonitrile + 0.1 % formic acid. The 10 min gradient started with 5 % solvent B, which was increased to 90 % by 5 min and further increased to 100 % solvent B by 7 min, before returning to 5 % solvent B at 7.1 min and held until 10 min (SI Fig. 10b).

### LC-MS/MS PFAS measurements per EPA Draft Method 1633

(Fig. 2d,f; SI Fig. 3d,f) Samples were analysed according to the EPA 3^rd^ Draft Method 1633^47^ and the corresponding Agilent application note^48^. In short, separation was performed using a ZORBAX Eclipse Plus column (C18, 2.1 × 100 mm, 1.8 μm; Agilent 959758-902) with a ZOBRAX Eclipse Plus (C18, 2.1 × 5 mm, 1.8 μm; Agilent 821725-901) guard column at 40 °C. The multisampler was kept at a temperature of 4 °C. The injection volume was 5 μl and the flow rate was 0.4 mL/min. The mobile phases consisted of A: 2 mM ammonium acetate in 95 % water + 5 % acetonitrile; B: acetonitrile. The 10 min gradient started with 2 % solvent B, which was increased to 95 % by 10 min, before returning to 2 % solvent B for a 2 min re-equilibration. LC-MS/MS analysis was performed on an Agilent 1290 Infinity II LC system coupled with an Agilent 6470 triple quadrupole (Agilent). The QQQ was operated in Dynamic MRM mode. The source parameters were as follows: gas temperature: 230 °C, gas flow: 6 L/min, nebulizer: 20 psi, sheath gas temperature: 355 °C, sheath gas flow: 10 L/min, VCap: 3500 V (positive) or 2500 V (negative), nozzle voltage: 2000 V (positive) or 0 V (negative). Pure standards (Agilent ITA-70) and labelled standards (Wellington Laboratories MPFAC-C-ES) were used for method development, compound identification and calibration (SI Figure 9c).

### LC-MS/MS for the analysis of PFNA

#### (Imperial College London; SI Fig. 3c)

Analysis was performed using a Shimdazu Nexera X2 LC-system coupled to a LCMS-8060 triple quadruple system with an electrospray ionisation source operated in negative ionisation mode (Shimadzu Corp., Kyoto, Japan). Separations were performed at 0.5 m/min using a Force C18 column (50 x 2.1 mm, 3 µm, Thames Restek, High Wycombe, UK) fitted with a matching guard column (Force C18, 5 x 2.1 mm, 5 µm) and a delay column (50 x 2.1 mm, 5 µm, Thames Restek) was installed between the mobile phase mixer and the autosampler. Sample order was randomised throughout the batch with an injection volume of 10 µl and the autosampler was held at 4 °C for the entire analysis. The elution program is as follows: an initial hold at 30 % mobile phase B (MPB, MeOH; MPA - 10 mM ammonium acetate (aq)) for 0.51 min, then an increase to 60 % MPB by 1.5 min, followed by another increase to 90 % MPB at 5 min, a hold at 95 % MPB from 6 to 7 min, followed by a 3-minute re-equilibration period at initial conditions. A 90:10 (MeOH:H2O) needle wash was used to rinse the outside of the needle before and after sample aspiration. Three transitions were monitored for both PFNA (463.05>419.00, 463.05>219.25, and 463.05>169.30) and M9PFNA (472.00>427.15, 472.00>223.10, and 472.00>172.25). Refer to Table 2 for the voltages and collision energies (optimised) used in each of the quadruples for each transition of PFNA and M9PFNA. Analysis of all extracts was accompanied by quality control samples (0, 1 and 10 µg/l in methanol, 1% ammonium hydroxide, 0.5 % acetic acid) and mobile phase blanks to account for instrumental contamination and to mitigate carryover.

**Table 2.** Transitions, voltages and collision energies for measured compounds.

| Compound | Precursor ion | Product ion | Q1 [V] | Collision Energy [V] | Q3 [V] | Polarity |
| --- | --- | --- | --- | --- | --- | --- |
| PFNA | 463.05 | 419 | 21 | 10 | 29 | negative |
| PFNA | 463.05 | 219.25 | 16 | 15 | 21 | negative |
| PFNA | 463.05 | 169.3 | 16 | 20 | 11 | negative |
| M9PFNA | 472 | 427.15 | 12 | 11 | 14 | negative |
| M9PFNA | 472 | 223.1 | 15 | 18 | 14 | negative |
| M9PFNA | 472 | 172.25 | 25 | 19 | 18 | negative |

### LC-MS/MS (QQQ) Metabolomics analysis (**Fig. 4d,e**; SI Fig. 7)

A deep-well plate containing 1180 µl PFNA in mGAM was inoculated with 10 µl second passage culture to create a final concentration of 20 µM PFNA. Samples were incubated at 37 °C for 24 h, after which 1 ml bacterial culture was transferred to a fresh tube and centrifuged at 4 °C and 14000 rpm for 3 min. Supernatant and pellet were collected and kept at −80 °C until extraction. Pellet samples and 100 µl of supernatant per sample were extracted with 800 µl of 1:1 methanol:water and vortexed; 200 µl chloroform was added to each sample and samples were incubated at −20 °C for 1 h for protein precipitation. Samples were then centrifuged at 4 °C and 14000 rpm for 2 min and the water phase was transferred to LCMS vials for analysis.

Amino acids and vitamins were quantified using liquid chromatography - tandem mass spectrometry as described previously^49^. On an Agilent 1290 Infinity II system, analytes were separated using hydrophilic interaction liquid chromatography (HILIC) with a Waters Acquity BEH Amide 1.7 µm, 2.1 mm X 100 mm column operated at 35°C. A binary buffer system of buffer A (50% acetonitrile, 10 mM ammonium formate, 0.176% formic acid) and buffer B (95:5:5 acetonitrile:methanol:water, 10 mM ammonium formate, 0.176% formic acid) was used at a constant flow rate of 0.9 ml/min and the following gradient: 0min:85%B, 0.7min:85%B, 2.55min:5%B, 2.9min:5%B, 2.91min:85%B, 3.5min:stoptime. The Agilent triple quadrupole 6470 instrument with JetStream ion source (AJS-ESI) was used in dynamic multiple reaction monitoring (dMRM) mode with a cycle time of 320 ms. The source parameters were as follows: Gas temp:325°C, Gas flow:10 l/min, Nebulizer:40psi, Sheath gas temp:350°C, Sheath gas flow:11 l/min, Capillary (positive):3500V, Capillary (negative):3500V, Nozzle voltage (positive):1000V, Nozzle voltage (negative):1000V. The injection volume was 0.25 µl. A serially diluted external calibration standard, blanks and a pooled QC sample was injected at regular intervals between samples. Data was analysed using MassHunter Workstation Quantitative Analysis for QQQ v10.1.

### LC-MS/MS data analysis

The Agilent MassHunter Qualitative Analysis 10.0 software was used to qualify the selected xenobiotic standards. TIC, EIC and EIC-fragment graphs were extracted for each compound. The Agilent MassHunter TOF Quantitative Analysis (Version 10.1) or Agilent MassHunter QQQ Quantitative Analysis software (Version 10.1) was used to quantify the xenobiotic compounds in each sample.

Calibration curves based on pure compound standards were used to estimate the concentrations of target compounds. Raw response values were used for concentration calculations for most analysis. Exceptions were the ‘Single strain-xenobiotic screen’, data represented in SI Fig. 2i, low concentration experiment per EPA Draft Method 1633, PFAS-mix experiment, and mouse fecal analysis, where the ratio of compound response to internal standard response was used. Data analysis was performed in RStudio Version 1.3.1093. The median of each sample group was compared to the median of the compound controls and an appropriate reduction of more than 20 % was chosen as cut-off, to ensure relevant reduction compared to the compound control distribution. In addition, statistical comparison was performed using t-test (two-sided), and p-values were FDR corrected (when a t-test was performed, the adjusted p-values are given in the respective figures). An adjusted p-value of less than 0.05 was considered significant.

In the ‘Single strain-xenobiotic screen’ bioaccumulation was defined as compound sequestration to at least 20 % from the supernatant but not from the whole culture sample, whereas biodegradation was defined as both supernatant and whole culture sample showing more than 20 % compound sequestration.

Concentration in samples extracted according to the EPA Draft Method 1633 were calculated based on the ratio with the labelled internal standard, which was spiked into the sample at a known concentration of 100 ng/l.

### PFAS-bacteria growth screens (**Fig. 1h**, SI Fig. 3b)

Plates (Corning 3795) containing 50 μl mGAM with 2x PFAS (2 % DMSO) concentration were prepared the evening prior and placed into the anaerobic chamber over night to ensure anaerobe conditions for inoculation. On the day of the screen each well was inoculated with 50 μl of a second passage culture to reach a starting OD_600_ of 0.05. Plates were sealed with a gas-permeable membrane (Breath-Easy, Merck, Cat# Z380059), which was additionally pierced with a syringe to prevent gas build-up. Plates were stacked without lids and incubated at 37 °C for 24 h in a stacker-incubator system (Biostack 4, Agilent BioTek) connected to a plate reader (Epoch 2, Agilent BioTek) to record the OD_600_ ^50^.

Growth curve analysis was performed in RStudio Version 1.3.1093. First, for each growth curve the minimum OD value was set to 0. Then the raw AUC was calculated for each well using ‘bayestestR’ package and area_under_curve() function. Further processing of growth curves was done by plate. AUC values were normalised by median AUC of all control wells (DMSO controls) on the respective plate to determine percent growth inhibition.

### Conventional ultrathin-section transmission electron microscopy (**Fig. 3a,b**; SI Fig. 4a-j)

On the day of the experiment each tube containing 2x concentration of PFAS was inoculated 1:1 with a second passage culture to reach a starting OD_600_ of 0.05. Samples were incubated at 37 °C for 24 h, after which bacterial cultures were spun down and the supernatant was removed. The bacteria were then fixed with a half Karnovsky fixative as 2.5% glutaraldehyde and 2 % paraformaldehyde in 0.1 M sodium cacodylate buffer (pH 7.4 with NaOH) for a few hours at room temperature. For conventional transmission electron microscopy^51^, the post-fixation was performed with a mixture of 1% osmium tetroxide and 1% potassium ferrocyanide in the cacodylate buffer. After stained en bloc with 5% aqueous uranyl acetate solution. The dehydration with a series of ethanol and the resin infiltration were completed for the plastic embedding in TER (TAAB Epoxy Resin). After the polymerization at 65 °C for a few days, the ultrathin-sections (∼60 nm) were cut by using an ultramicrotome (Leica EM UCT/UC7/Artos-3D, Vienna Austria), mounted on formvar-carbon films on EM copper grids, and stained with lead citrate. The bacterial ultrastructure was observed using FEI Talos F200C 200kV transmission electron microscope (Thermo Fischer Scientific, Einthoven Netherlands) with Ceta-16M CMOS-based camera (4kx4k pixels under 16bit dynamic range) and JEM-1400 Flash TMP (JEOL Ltd., Tokyo Japan) with TVIPS TemCam-XF416 CMOS (Tietz Video and Image Processing Systems GmbH, Germany) as described previously^52^.

### Automated TEM image sectioning and analysis (**Fig. 3c-f**; SI Fig. 4k,l)

Electron microscopy images of were converted to 8-bit format using FIJI^53^. Only images with the same magnification level were analysed. The segmentation and quantification of these images were performed using CellProfiler 4.2.6 (Stirling, DR, BMC Bioinformatics, 2021^54^). Individual cells were segmented as primary objects based on their intensity and size after the application of a Gaussian blur filter. Within each cell, condensates were also segmented based on intensity and size. Mean pixel intensity and number of condensates were quantified for each cell and their corresponding condensates and the results were exported as CSV files. Cells touching the edges of the images were not considered for this quantification. Images showing the segmented areas were saved as TIFF files. All sectioned cells and aggregates were manually checked and wrong sectioned cells and cells that were identified in duplicate were excluded from further analysis.

### Cryogenic FIB-SIMS imaging

Cryogenic focused ion beam secondary ion mass spectrometry (FIB-SIMS) imaging^55^ was performed on a focused ion beam scanning electron microscope (FIB-SEM, Zeiss Crossbeam 550) equipped with a time-of-flight mass spectrometer (Tofwerk). During FIB-SIMS imaging, the sample is scanned by a Gallium focused ion beam, which ablates material at each pixel. The secondary ions resulting from the interaction of ion beam and sample are extracted at each scanning position and are analysed by time-of-flight secondary electron mass spectrometry (TOF-SIMS). The method hence allows a pixel-by-pixel visualisation of the sample’s chemical composition. The simultaneous detection of secondary electrons yields spatial images of the sample (FIB images, Fig. 3g). As material is continuously removed during the imaging process, repeated scanning of a given sample area provides three-dimensional information regarding both spatial features and chemical composition of the sample, thus creating a volume with a mass spectrum for each voxel. This not only allows this method to show the lateral localisation of elements of interest, but also provides depth information: each two-dimensional image resulting from a full scan of the observed area can be considered as a slice of the sample (frame) within a Z-stack (Fig. 3) while the distribution along the Z-axis can be extracted as X-Z slices. To retain a near-native state of the sample, *E. coli ΔtolC* cells were plunge-frozen in liquid ethane using a Vitrobot Mark IV (ThermoFisher) after applying 2.5 µl of sample to a cryoEM grid (Quantifoil Cu/Rh R3.5/1, UltrAuFoil R 1.2/1.3) as described previously^56^.

### Proteomics analysis (**Fig. 4a**; SI Fig. 6)

A deep-well plate containing 1180 µl PFNA in mGAM was inoculated with 10 µl second passage culture to create a final concentration of 20 µM PFNA. Samples were incubated at 37 °C for 24 h, after which bacterial cultures were spun down and the supernatant was removed.

The bacterial cell pellets were lysed in 100 mM Triethyl ammonium bicarbonate (TEAB) containing 0.1% RapiGest surfactant by sonication on ice (50J x5, 30% Amplitude) after denaturing by two rounds of incubation at 80 °C for 10 min with intermittent cooling. The solubilized protein content was estimated using Pierce 660nm assay as per manufacturer’s instructions. 50 µg of protein from each sample was reduced with 4 mM dithiothreitol (DTT), alkylated with 14 mM iodoacetamide (IAA), and further digested with trypsin at 1:50 protease:protein ratio for 16h at 37 °C. The peptide content was estimated at 1:10 dilution using Pierce quantitative colorimetric assay as per manufacturer’s instructions. Equal amounts of peptides from respective bacterial samples were labelled with TMT in a randomized fashion to ensure the reporter ion isotopic distribution is spread across the replicates following manufacturer’s guidelines (Thermo Fisher Scientific). Equal volume of the respective TMT labelled peptides were pooled together and desalted to remove unbound labels. The desalted peptides were further subjected to fractionation based on their reverse phase chromatographic properties under basic pH using an analytical HPLC. The 12 concatenated fractions collected were vacuum dried, resuspended in 0.1% TFA containing 3% acetonitrile (ACN), and taken for LC-MS/MS analysis.

LC-MS/MS analysis was carried out in RTS-SPS-MS3 mode for TMT reporter ion quantification. An equal amount of predetermined sample load from each of the 12 fractions were analysed using Orbitrap Eclipse™ mass spectrometer with an Ultimate 3000 RSLC™ nano system coupled in-line. The peptides were loaded onto the trapping column (Thermo Fisher Scientific, PepMap100, C18, 300 μm X 5 mm), using partial loop injection, washed for three minutes at a flow rate of 15 μl/min with 0.1% formic acid(FA). The peptides were resolved on an analytical column (Easy-Spray C18 75 µm x 500 mm; 2 µm particle size) at a flow rate of 300 nL/min using a gradient where %B is raised from 3-25 over 140 minutes and then to 40%for additional 13 minutes. The column was washed in 90% B for 12 minutes and re-equilibrated in 3%B for 15 minutes before next injection. 0.1% formic acid (FA) in water was used as mobile phase A and 0.1% FA in 80% ACN was used as mobile phase B. Data was acquired using three FAIMS compensation voltages (−45v, −60v and −75v) For each FAIMS experiment with a maximum cycle time of 2s, mass spectra acquisition was carried out in RTS-SPS-MS3 mode. In this mode, full-scan MS was acquired in the mass range of 415-1500 m/z at 120,000 resolution with a maximum ion injection time of 30ms (AGC target 2e5 ions). Precursors selected for MS/MS were isolated using an isolation width of 0.7 m/z and fragmented by HCD using collision energy of 32%. MS/MS scan was performed on the ion trap at turbo scan rate (AGC 5e4 ions) with a maximum ion injection time of 22 ms. To avoid repeated selection of peptides for MSMS the program used a 40 second dynamic exclusion window. Real time search parameters were set as follows using respective fasta databases; Uniprot *Bacteriodes uniformis* database, UP000014212 (unreviewed db downloaded on 20230621, contains 3957 seq), Uniprot *Odoribacter splanchnicus* database, UP000006657 (unreviewed db downloaded on 20230621, contains 3479 seq) Uniprot *E.coli* K12 database, UP000000625 (reviewed db downloaded on 20230621, contains 4362 seq). Protease: trypsin, static modifications: cysteine carbamidomethylation (+57.0215) and TMT (+229.163) or TMTPro (+304.207) on lysine and N-terminus, Variable modification: Methioninie oxidation (+15.99491). One missed cleavage was allowed, and FDR filtering was enabled. Only 5 peptides per protein were allowed per basic reverse phase fraction (using the close-out option) and maximum search time for RTS was set to 35 ms. Ten of the most abundant peptide fragments were selected for SPS-MS3 and MS3 spectrum was acquired at 120,000 resolution on the mass range 100-500 m/z with an AGC target of 500% and a maximum injection time of 246 ms.

Data was processed using Proteome Discoverer v3.0 using the protein databases mentioned above. The spectra identification was performed with the following parameters: MS accuracy, 10 ppm; MS/MS accuracy, 0.6 Da for spectra acquired in Ion trap mass analyser; missed cleavage, up to 2; fixed modifications, carbamidomethylation of cysteine and TMT/TMTpro on lysine and on peptide N-terminus; variable modifications, oxidation of methionine. An intensity-based rescoring of PSMs identified in SequestHT node was performed by Inferys prior to false discovery rate estimation using Percolator. Only rank 1 peptide identifications with high confidence (FDR<1%) were accepted for identification and quantification reporting. Quantification was carried out on the S/N values of reporter ions for the identified peptides that are unique to the proteins. The protein abundance was enumerated by summing the abundances of the connected peptide groups matching to respective proteins. Data normalization was carried out using the TMT channel with the highest total protein abundance as the reference channel. The normalized abundance values are reported for comparison across datasets.

### Thermal proteome profiling (TPP) (**Fig. 4b,c**)

TPP was performed as previously described^23,57^. In brief, cells were grown anaerobically at 37 °C for 48 h. Cells were then washed twice with anaerobic PBS. For the live cell experiments, cells were treated with 8 different concentrations of anaerobic PFNA (2-fold dilutions starting at 10 µM including DMSO control) for 30 min at 37 °C in the anaerobic chamber. Aliquots of treated cells were then incubated in 10 different temperatures for 3 min. Cells were resuspended in lysis buffer (50 µg/ml lysozyme, 1mM MgCl2, cOmplete protease inhibitors, 0.25 U/µl benzonase in PBS) with 0.8% NP-40 and lysed with 5 freeze-thaw cycles. Aggregates were then removed and soluble proteins were prepared for mass spec. For the lysate experiments cells were lysed as above but without the presence of NP-40. NP-40 was added after the heat treatment before the aggregate removal.

Proteins were digested as previously described^58,59^. Eluted peptides were labelled with TMT16plex, pooled (all treatments of each two adjacent temperatures per experiment) and pre-fractionated into 6 fractions under high pH conditions. Samples were then analyzed with liquid chromatography coupled to tandem mass spectrometry, as previously described^57^.

All raw files were converted to mzmL format using MSConvert from Proteowizard, using peak picking from the vendor algorithm. Files were then searched using MSFragger v3.7^60^ in Fragpipe v19.0 against the uniprot proteome UP000000625 containing common contaminants and reversed sequences.

Data analysis was performed using R. Only proteins with at least two identified peptides were kept for the analysis and outlier conditions were removed altogether. For the correlation analysis, per experiment, protein intensities across all concentrations within each temperature were normalized using variance stabilization normalization^61^ and the cumulative fold change across temperatures and concentrations per protein was calculated. For the hit calling the data were preprocessed using the TPP package^40^ and analysed using the TPP2D package^62^ with an alpha of 0.1.

### Adaptive laboratory evolution (SI Fig. 5)

*B. uniformis*, *B. thetaiomicron*, *O. splanchnicus*, *P. merdae* and *E. coli* BW25113 Δ*tolC* were evolved for 20 days in 500 μM PFHpA, 500 μM PFOA, 250 μM PFNA, and 125 μM PFDA. DMSO was used as a control. For each PFAS compound 4 and for the DMSO controls 8 replicate lineages were evolved in parallel. 2 ml deep-well stock plates containing a 100x stock of each PFAS in DMSO were prepared prior to the experiment and stored at −80 °C until use. On day 0 each well was inoculated with a second passage culture to reach a starting OD_600_ of 0.05. On each of the following days 50 μl of grown culture was transferred to a fresh compound plate containing PFAS/DMSO in mGAM. Every 5 days the growth of the strains in presence of PFAS was measured by transferring 100 μl starting culture to a clear-bottom plate and measured and analysed it as described in the section ‘PFAS-bacteria growth screens’. On day 20 glycerol stocks were prepared from each lineage and stored at −80 °C.

#### Genome sequencing

Genomic DNA of B. uniformis after evolving in PFNA, PFOA and DMSO control was extracted with the QIAamp PowerFecal Pro DNA kit (Qiagen; 51804) and quantified using Qubit 1X dsDNA HS Assay Kits (Q33230). On average, 74.38 ng/µl of DNA was obtained and sequenced by Illumina NovaSeq 6000 with a library insert size of 350 bp, resulting in 6.50 to 9.95 million paired reads (mean = 8.68 million) with a length of 150 bp. Sequencing reads were then filtered with fastp (v 0.32.2)^63^ using default parameters. We called the variants in evolved strains using snippy pipeline (https://github.com/tseemann/snippy). Filtered reads from PFNA, PFOA and DMSO evolved population were aligned to the reference genome with BWA and variants were called by freebayes^64^ using parameters “--mincov 10 --basequal 30 - -fbopt ‘-C 3 -p 1 -F 0.01’” defined through snippy. Afterward, those variants appearing in both PFNA/PFOA and DMSO control samples were removed.

### Mouse experiments (**Fig. 5**; SI Fig. 8)

Animal experiments were approved by the local authorities (Regierungspräsidium Tübingen, H 02/20 G). Germfree C57BL/6J mice were bred in house (Gnotobiotic Mouse Facility, Tübingen). Mice were housed under germfree conditions in flexible film isolators (Zoonlab) and transferred to the Isocage P system (Tecniplast) to perform the experiments. Mice were supplied with autoclaved drinking water and γ-irradiated maintenance chow for mice (Altromin) ad libitum. Female and male mice between 5 – 6 weeks were used and animals were randomly assigned to experimental groups. Mice were kept in groups of 3 mice per cage during the experiment. All animals were scored daily for their health status.

#### Preparation and inoculation of the Com20 and high-vs. low-accumulating bacterial community

The Com20 and high- and low-accumulating communities (SI Table 37) were prepared under anaerobic conditions (Coy Laboratory Products Inc., 2 % H_2_, 12 % CO_2_, rest N_2_). Consumables, glassware and media were pre-reduced at least 2 days before inoculation of bacteria. Each strain was grown in monoculture overnight in 5 ml of their respective growth medium at 37 °C. The next day, bacteria were sub-cultured (1:100) in 5 ml fresh medium and incubated for 16 h at 37 °C, except *Eggerthella lenta*, which was grown for 2 days. Optical density (OD) at 578 nm was determined and bacteria were mixed together in equal ratios to a total OD of 0.5 in a final volume of 10 ml. After adding 2.5 ml 50 % glycerol (with a few crystals of palladium black (Sigma-Aldrich)), 200 µl aliquots were prepared in glass vials (2 ml, Supelco, Ref. 29056-U) and directly frozen at −80 °C. Frozen vials were used within 3 months.

Inoculation of germfree mice was performed according to Grießhammer et al.^46^. In short, cages were transferred to an ISOcage Biosafety Station (IBS) (Tecniplast) through a 2 % Virkon S disinfectant solution (Lanxess) dipping bath. Glycerol stocks of the frozen communities (one per mouse) were kept on dry ice before being thawed during transfer into the IBS. Mixtures were used directly after thawing with a minimal exposure time to oxygen of maximum 3 min. Mice were inoculated by oral gavage (50 µl) and inoculation was repeated after 48 h using the same protocol. The germfree control group was left untreated. The IBS was sterilized with 3 % perchloracetic acid (Wofasteril, Kesla Hygiene AG).

#### Fecal sample collection

10 days after the second inoculation with Com20, high-or low-accumulating communities, mice were orally gavaged with PFNA (10 mg/kg or 0.1 mg/kg in 25 % DMSO) in a volume of 50 µl. Fresh fecal samples were collected before treatment, 3 h, 1 day and 2 days after treatment in sterile 1.5 ml Eppendorf-tubes and immediately frozen at −80 °C. On day 3 after treatment, mice were euthanized by CO_2_ and cervical dislocation, dissected and intestinal contents were taken from colon and the small intestine and collected in the same way.

#### 16S rRNA gene sequencing of fecal samples

DNA extraction from fecal samples was performed in house using MagMAX^TM^ Microbiome Ultra Nucleic Acid Isolation Kit in bead tubes (Thermo Fisher Scientific, A42358) and the KingFisher^TM^ Flex (MAN0018071) according to the manufacturer’s instructions. After, DNA integrity was verified using agarose gel electrophoresis and the DNA concentration was determined using Qubit^TM^ dsDNA BR assay kit (Thermo Fisher Scientific, Q32853) in combination with the Varioskan LUX plate reader (thermoscientific).

Two-step PCR protocol was used to prepare 16S amplicons. A single amplicon was generated by using V4 primer pairs (515F / 806R) for the first mouse experiment (10 mg/kg PFNA, Com20 vs. GF) and V4 and V5 primer pairs (515F / 926R) for second mouse experiment (10 mg/kg PFNA, high-vs. low-accumulating community). In the first step PCR, primers with overhang adapter sequences were added; afterwards in the 2nd PCR Illumina sequencing adapters and dual-index barcodes were added to the amplicon target for pooling. Pooled samples are sequenced on Illumina MiSeq instrument using 2×250 PE protocol at the Genomics Core Facility (EMBL Heidelberg).

In total, 1,086,628 pair-end reads for the first mouse experiment and 6,531,459 for the second mouse experiment (average 10,061 pairs per sample in the first experiment and 56,306 pairs per sample in the second experiment) with 250 bp were generated. Raw sequencing reads were truncated and filtered using the DADA2 pipeline (v1.26.0)^65^ with the following parameters: “truncLen=c(230,210), maxN=0, maxEE=c(2,2), truncQ=2, rm.phix=TRUE”. Afterwards, error rates were learned from filtered reads, and corrected reads were merged as ASVs. A self-defined reference was used for alignments of ASVs and calculation of relative abundances.

### Data analysis and replicates

All data analysis was performed using open-source packages accessed from RStudio (Version 1.3.1093). The Agilent MassHunter TOF Quantitative Analysis (Version 10.1) or Agilent MassHunter QQQ Quantitative Analysis software (Version 10.1) was used to quantify the xenobiotic compounds in each sample. All t-tests are two-sided. All p-values are FDR corrected. Biological replicates refer to different inoculation cultures, while technical replicates refer to experiments starting with the same inoculation culture. None of the data points correspond to repeated measurements of the same sample. Fold changes refer to the ratio of medians.

### Data and code availability

All data is included in the supplementary information. All electron microscopy images are available at EBI Bioimage archive (TMP_1715003294892). The raw mass-spectrometry data will be available at EBI MetaboLights repository (MTBLS9756 and MTBLS9745). The raw TPP data have been deposited to the ProteomeXchange Consortium via the PRIDE^66^ partner repository with the dataset identifier PXD050999. Raw sequencing reads of the evolved *B. uniformis* and 16S rRNA genotyping were uploaded to the European Nucleotide Archive (ID PRJEB72794, PRJEB75767). No custom algorithms used for data analysis; the code used is available upon request.

