## Supplementary figures for "Extensive PFAS accumulation by human gut bacteria"

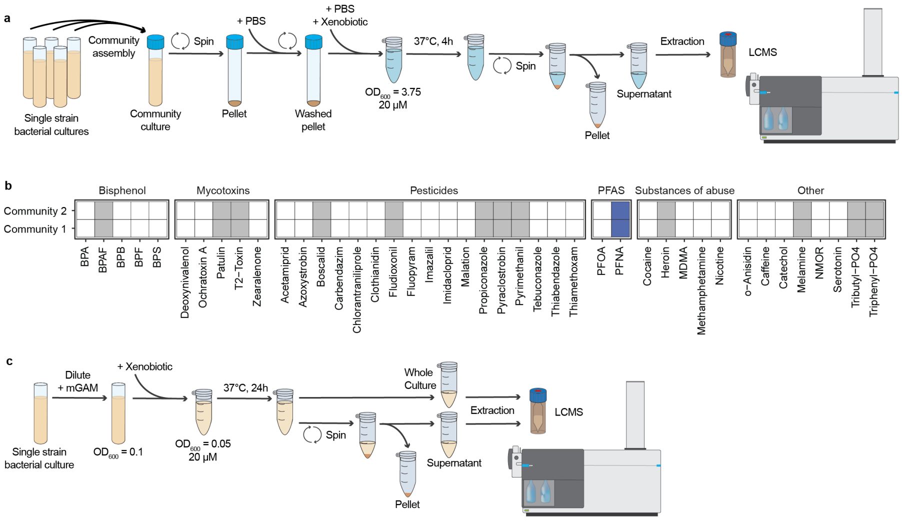


**Supplementary Figure 1. Method setup and results from the community-screen. a.** Method workflow for the artificial community experiment. Each community consisted of 10 bacteria (community 1: *Bacteroides caccae, Bacteroides dorei, Bacteroides thetaiomicron, Bacteroides uniformis, Bacteroides vulgatus, Colinsella aerofaciens, Coprococcus comes, Eubacterium rectale, Parabacteroides merdae, Roseburia intestinalis;* community 2: *Akkermansia muciniphila, Bacteroides clarus, Bacteroides stercoris, Clostridium difficile, Eggerthella lenta, Eubacterium eligens, Fusobacterium nucleatum subsp. animalis, Odoribacter splanchnicus, Parabacteroides distastonis, Ruminococcus bromii*). **b.** 13 out of 42 tested pollutants were sequestered by at least one synthetic gut bacterial community. Coloured squares denote sequestration, i.e., >20 % reduction from the bacteria-free supernatant. PFAS are marked in blue. n = 6 (3 biological and 2 technical replicates) (SI Table 3). **c.** Method workflow for the single strain experiment.

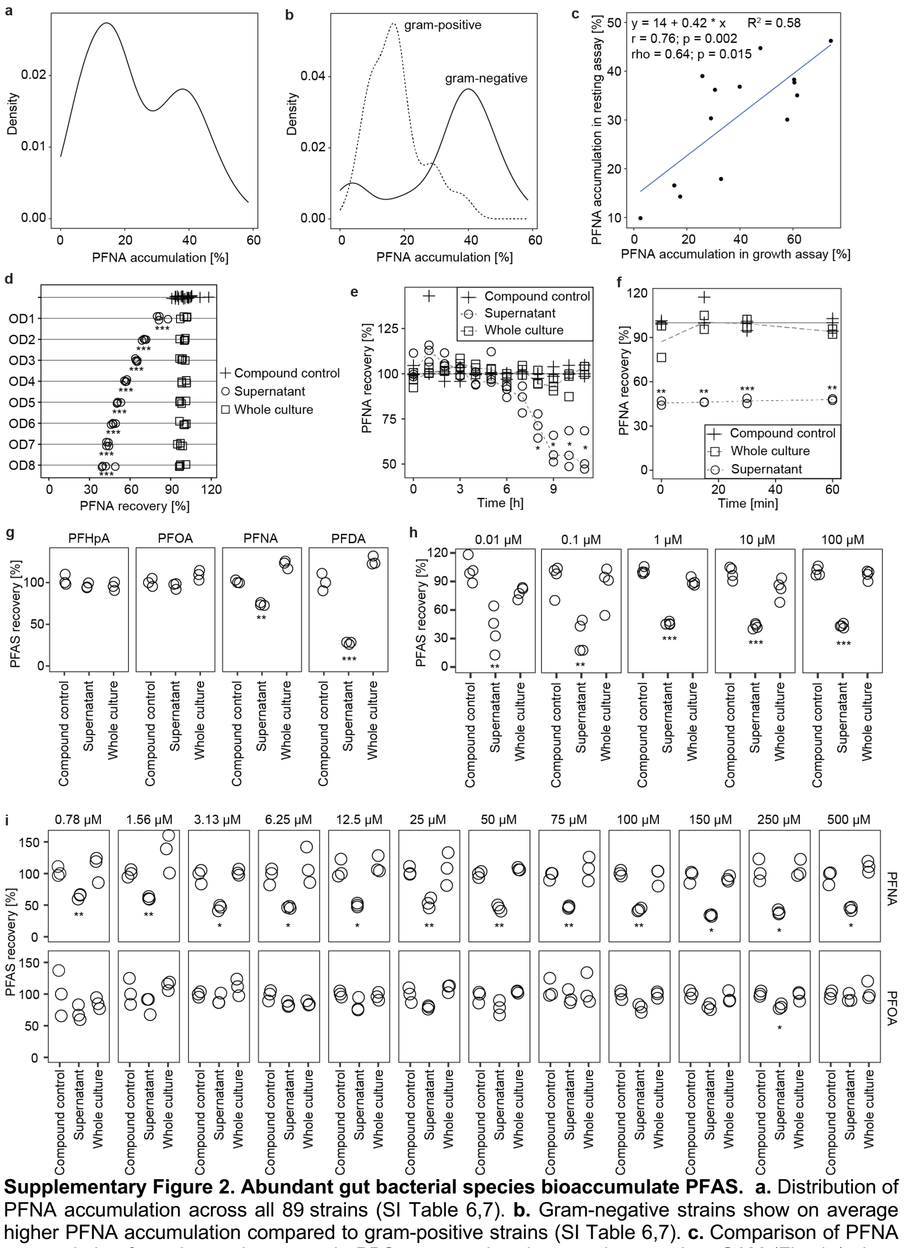

**Supplementary Figure 2. Abundant gut bacterial species bioaccumulate PFAS. a.** Distribution of PFNA accumulation across all 89 strains (SI Table 6,7). **b.** Gram-negative strains show on average higher PFNA accumulation compared to gram-positive strains (SI Table 6,7). **c.** Comparison of PFNA accumulation from the resting assay in PBS compared to the growth assay in mGAM (Fig. 1a) show strong positive correlation (Pearson rank correlation: r = 0.76, p-value = 0.002; Spearman rank correlation: ρ = 0.64, p-value = 0.015). **d**. Data from Fig. 1c including compound control and whole culture samples. PFNA depletion by *B. uniformis* cultures of varying $OD_{600}$ in PBS buffer. All ODs show significant PFNA accumulation compared to the compound control. $OD_{600}$ = 1-8; 20 μM initial PFNA concentration; *** p < 0.001; n = 4 technical replicates (SI Table 8). **e.** Data from Fig. 1d including compound control and whole culture samples. Kinetics of PFNA depletion during *B. uniformis* growth starting with low cell density. A significant amount of PFNA was sequestered from the media after 8 h of growth and onwards. initial $OD_{600}$ = 0.05; 20 µM initial PFNA concentration; * p<0.05 and >20% PFNA sequestration from the media; n = 3 biological replicates (SI Table 9,10). **f.** Kinetics of PFNA depletion by *B. uniformis* when starting with high cell density in mGAM ($OD_{600}$ = 4). Bioaccumulation of ca. 50 % PFNA happens within the time frame of sample collection (ca. 5 min). ** p-value < 0.01; *** p-value < 0.001; n = 2 biological replicates (SI Table 12). **g.** Data from Fig. 1g including compound control and whole culture samples. Bioaccumulation of PFAS compounds with varying chain length by *B. uniformis*. PFNA and PFDA are significantly accumulated compared to the compound control. $OD_{600}$ = 3.75; 20 µM initial concentration for all compounds; ** p-value < 0.01; *** p-value < 0.001; n = 3 technical replicates (SI Table 14). **h.** Data from Fig. 1f including compound control and whole culture samples. PFNA is bioaccumulated by *B. uniformis* grown in mGAM at a range of initial concentrations compared to the compound control. initial $OD_{600}$ = 0.05; initial PFNA concentrations = 0.01 to 100 µM; ** p-value < 0.01; *** p-value < 0.001; n = 4 technical replicates (SI Table 13). **i.** PFNA and PFOA are bioaccumulated by *B. uniformis* in PBS at a range of initial concentrations. $OD_{600}$ = 3.75; initial PFAS concentrations = 0.78 to 500 µM; ** p-value < 0.01; *** p-value < 0.001; n = 3 technical replicates (SI Table 15).

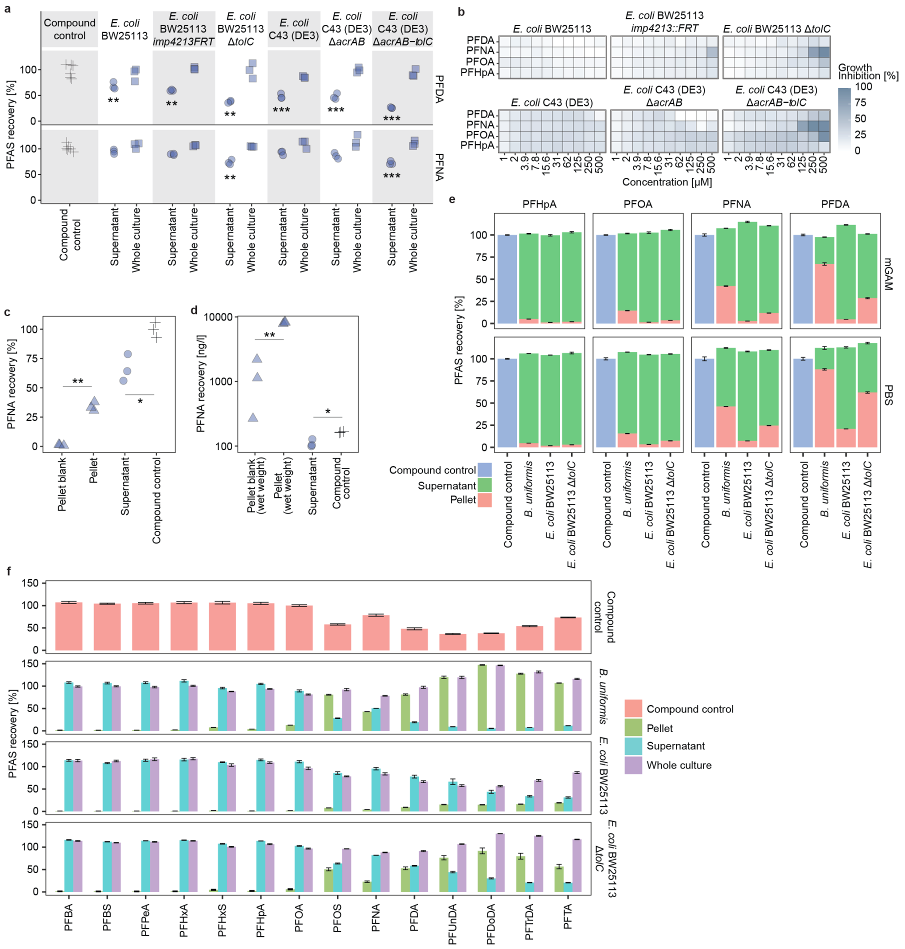


**Supplementary Figure 3. a.** Data from Fig. 2c including compound control and whole culture samples. Accumulation of PFDA and PFNA by wild-type *E. coli* strains and corresponding efflux mutants. $OD_{600}$ = 3.75; exposure concentration = 20 µM; ** p-value < 0.01 and >20 % sequestration compared to the compound control; *** p-value < 0.001 and >20 % sequestration compared to the compound control; n = 3 technical replicates (SI Table 19). **b.** Efflux mutants show increased PFAS sensitivity. n = 3 technical replicates (SI Table 16,17). **c.** Independent measurement of samples from Fig. 2d at Imperial College London supports PFNA accumulation by *B. uniformis* at 160 ng/l exposure concentration. * p-value < 0.05; ** p-value < 0.01; n = 3 biological replicates (SI Table 22). **d.** Data from Fig. 2d displayed in a different way: based on wet pellet weight, concentration within the bacterial pellet was a median of circa 8200 ng/l, which is a 50-fold increase compared to the exposure concentration. * p-value < 0.05; ** p-value < 0.01; n = 3 biological replicates (SI Table 20,21). **e.** Data related to Fig. 2e: PFAS recovery from pellet and supernatant after exposure to 5 µM PFAS in mGAM (initial $OD_{600}$ = 0.05, 24 h incubation) or PBS ($OD_{600}$ = 3.75, 4 h incubation). n = 3 technical replicates (SI Table 24). **f.** Data from Fig. 2f including compound control, whole culture and supernatant samples. PFAS recovery after 1 h exposure in PBS ($OD_{600}$ = 3.75, PFAS mix of 14 compounds each at a concentration of 1 mg/l). For PFAS compounds with a chain length of more than ten carbon atoms, the solubility is limited at this concentration; the low accumulation in *E. coli* BW25113 wild-type pellets acts as a biological control in this case. n = 3 technical replicates (SI Table 26).

**Commented [AL1]:** Error bars show standard error

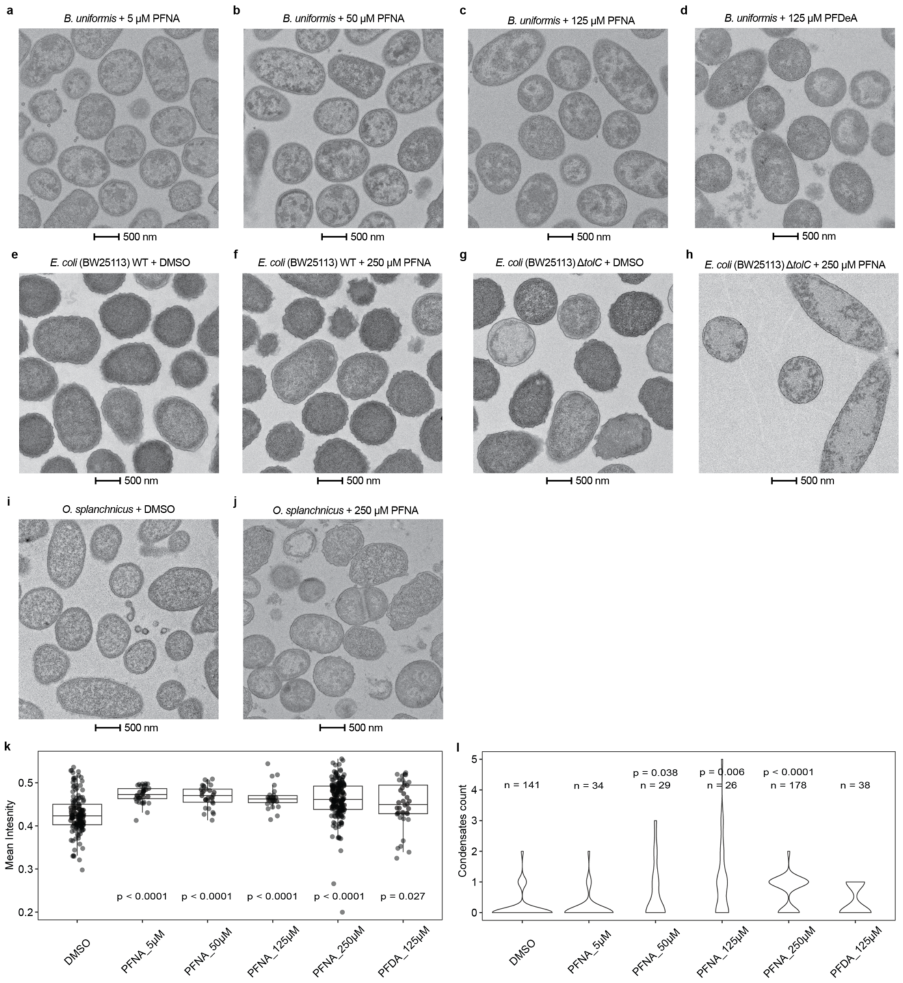


**Supplementary Figure 4. Morphological data support intra-cellular accumulation of PFAS. a-d.** TEM of *B. uniformis* cells grown in mGAM + 5 µM PFNA (**a**), 50 µM PFNA (**b**), 125 µM PFNA (**c**), or 125 µM PFDA (**d**) for 24 h. **e,f.** TEM of *E. coli.* BW25113 wild-type cells grown in mGAM + DMSO **(e)** or 250 µM PFNA **(f)** for 24 h. **g,h.** TEM of *E. coli.* BW25113 Δ*tolC* cells grown in mGAM + DMSO **(g)** or 250 µM PFNA **(h)** for 24 h. **i,j.** TEM of *O. splanchnicus* cells grown in mGAM + DMSO **(i)** or 250 µM PFNA **(j)** for 24 h. **k.** *B. uniformis* data from Fig. 3e including data for 5, 50, 125 µM PFNA and 125 µM PFDA. Mean intensity of bacterial cells show significant differences between DMSO and PFNA or PFDA treated cells (SI Table 27). **l.** *B. uniformis* data from Fig. 3f including data for 5, 50, 125 µM PFNA and 125 µM PFDA. Condensate count per bacterial cell show significant increase in condensates for *B. uniformis* exposed to 50, 125 and 250 µM PFNA, supporting morphological changes through PFNA exposure (SI Table 27).

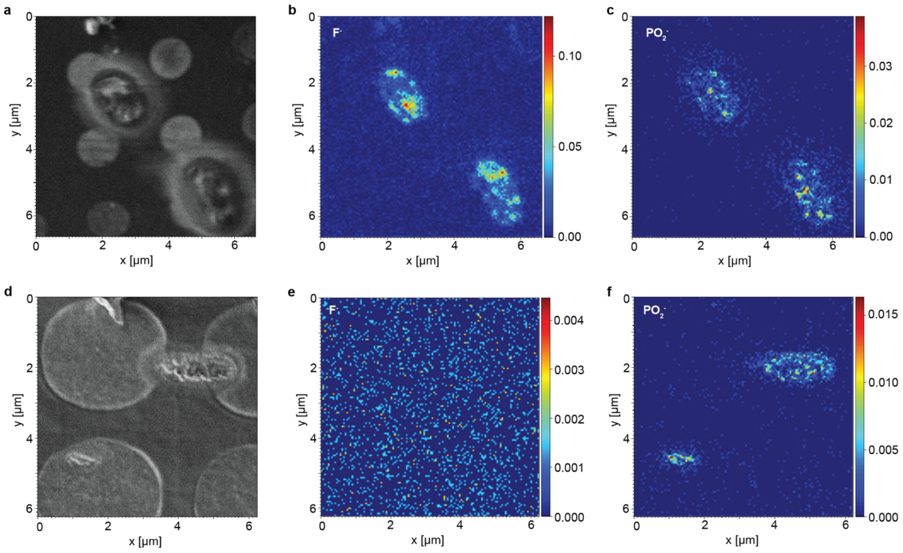


**Supplementary Figure 5. Cryogenic FIB-SIMS of *E. coli* Δ*tolC* exposed to 250 µM PFNA show intra-cellular accumulation of PFAS compared to the DMSO control.** During cryogenic FIB-SIMS imaging, the vitrified sample is scanned by a Gallium focused ion beam (FIB) and time-of-flight secondary electron mass spectrometry (ToF-SIMS) is performed on the secondary ions created by the interaction of ion beam and sample at each scanning position. Thus, the chemical composition of the sample can be visualised pixel-by-pixel. **a-c.** *E. coli* Δ*tolC* exposed to 250 µM PFNA. **d-f.** *E. coli* Δ*tolC* exposed to DMSO control. For both cases, a spatial image of a cell resulting from the secondary electrons created during FIB imaging is shown on the left (**a,d**). Please note that because the specimens were frozen on a holey cryo-EM grid, round holes in the film are additionally observed, in addition to the elongated cells. On the right (**b,c,e,f**), ToF-SIMS images of the same cell are shown for mass-to-charge ratios 19 and 63, corresponding to $F^-$ and $PO_2^-$, respectively. The colour bars indicate ion count, i.e., regions shown in red are those where the most ions of the particular species were detected. $F^-$ signal indicates the presence of PFNA, $PO_2^-$ was chosen as a comparison as it occurs in most bacterial cells and hence is a reliable marker for cells in negative SIMS imaging mode. In *E. coli* Δ*tolC* cells exposed to PFNA, a strong $F^-$ signal is observed within the cell (**b**), which is not the case for the DMSO treated cells (**e**), in which a faint background signal can only be observed when adjusting the colour scale by several orders of magnitude.

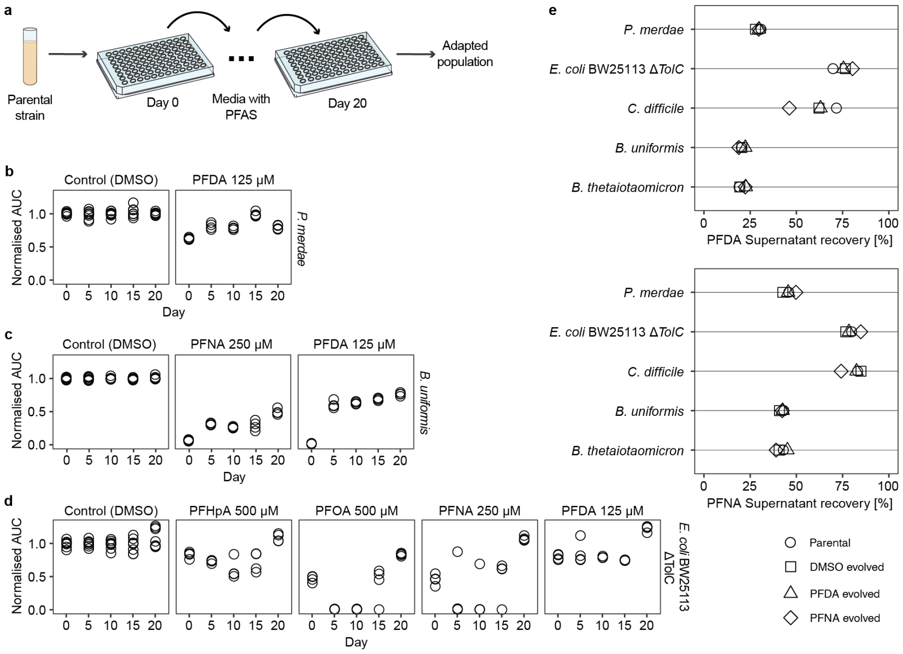


**Supplementary Figure 6. PFAS tolerance and bioaccumulation following adaptive laboratory evolution. a.** Five gut bacterial species were evolved through serial passaging in growth medium containing one of four PFAS compounds (500 µM PFHpA, 500 µM PFOA, 250 µM PFNA, 125 µM PFDA) over 20 days. **b.** Improved growth of adapted *P. merdae* in presence of 125 µM PFDA at days 5, 10, 15 and 20 compared to day 0 (day 20: 1.3-fold change, p-value = 0.001). n = 4 independent populations per compound (SI Table 28,29). **c.** Improved growth of adapted *B. uniformis* population in presence of 250 µM PFNA and 125 µM PFDA after 5, 10, 15 and 20 days compared to day 0 (day20 PFNA 7-fold change, p-value = 0.0004; day 20 PFDA 46-fold change, p-value = 0.00003). n = 4 independent populations per compound (SI Table 28,29). **d.** Improved growth of adapted *E. coli* BW25113 *ΔTolC* population in presence of 500 µM PFHpA, 500 µM PFOA, 250 µM PFNA and 125 µM PFDA at day 20 compared to day 0 (PFHpA 1.3-fold change, p-value = 0.002; PFOA 1.7-fold change, p-value = 0.0006; PFNA 2.3-fold change, p-value = 0.0006; PFDA 1.6-fold change, p-value = 0.00005). n = 4 independent populations per compound (SI Table 28,29). **e.** Adapted populations retain PFDA and PFNA bioaccumulation capability. Mean of n = 4 independent populations per compound (SI Table 30).

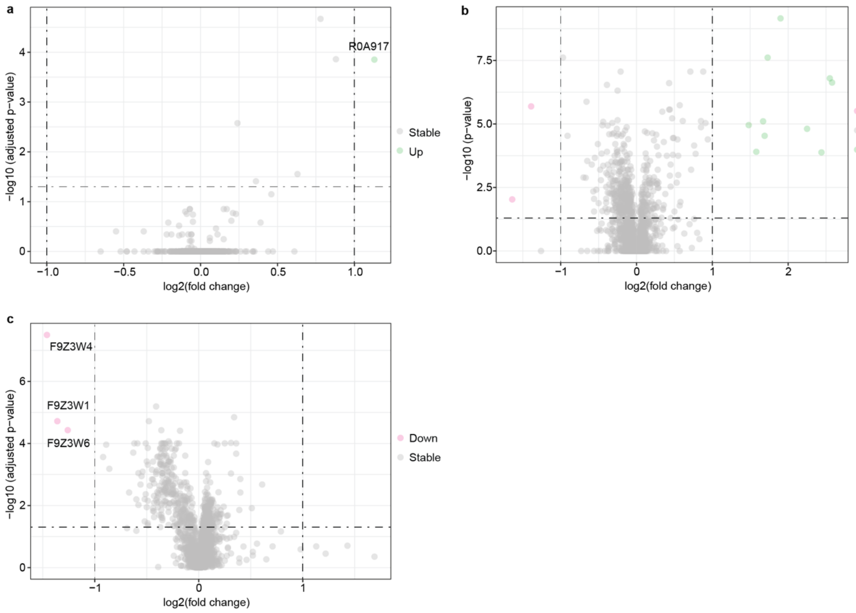


**Supplementary Figure 7. Proteomics results show effects of PFNA on abundance of proteins. a-c.** Results showing the proteins that are differentially abundant between (**a**) *E. coli* BW25113, (**b**) *E. coli* BW25113 *ΔTolC,* and (**c**) *O. splanchnicus* treated with 20 µM PFNA in comparison to DMSO. The green and the blue dots mark proteins with a log2 abundance ratio >1 or <-1 (i.e., two-fold increase or decrease) and a multiple-testing corrected p-value of less than 0.05. n = 6 biological replicates (SI Table 32).

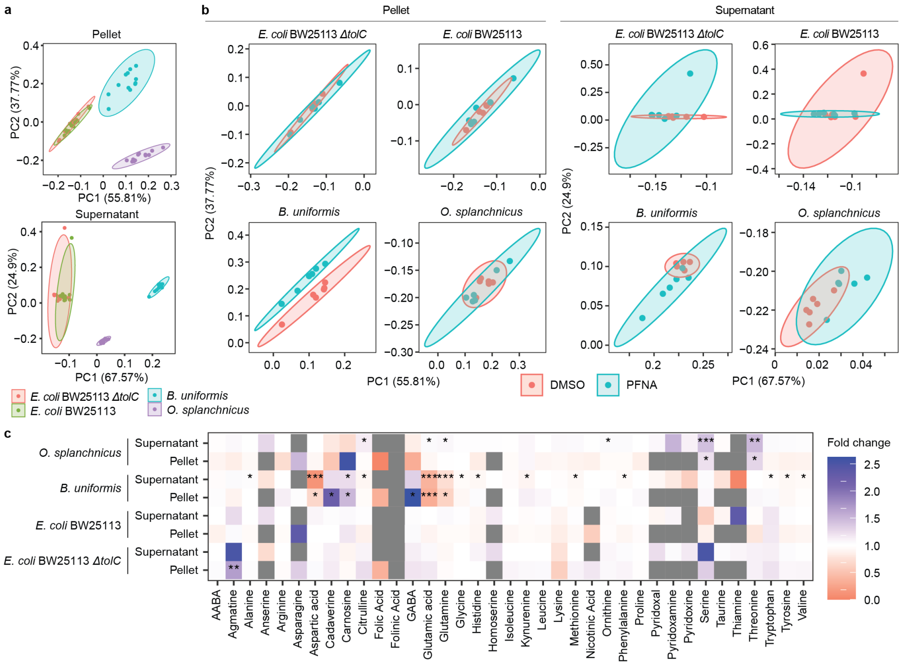


**Supplementary Figure 8. Metabolomics results show effects of PFNA on abundance of 37 metabolites in pellet and supernatant samples. a.** PCA analysis shows clear distinction between bacterial strains, with E. coli wild-type and TolC mutant overlapping. n = 6 biological replicates (SI Table 35). **b.** PCA analysis of pellet and supernatant samples shows clear distinction of PFNA and DMSO treated cells only for *B. uniformis* pellet samples (*B. uniformis* pellet PCA is a replicate of Fig.4d, added for completion). n = 6 biological replicates (SI Table 35). **c.** Metabolomics analysis show change in intra- and extracellular metabolites. Fold change of analysed metabolites in supernatant and pellet samples of 20 µM PFNA exposed cells compared to control (DMSO) samples. Significance: * p<0.05, ** p<0.01, *** p<0.001. n = 6 biological replicates (SI Table 36).

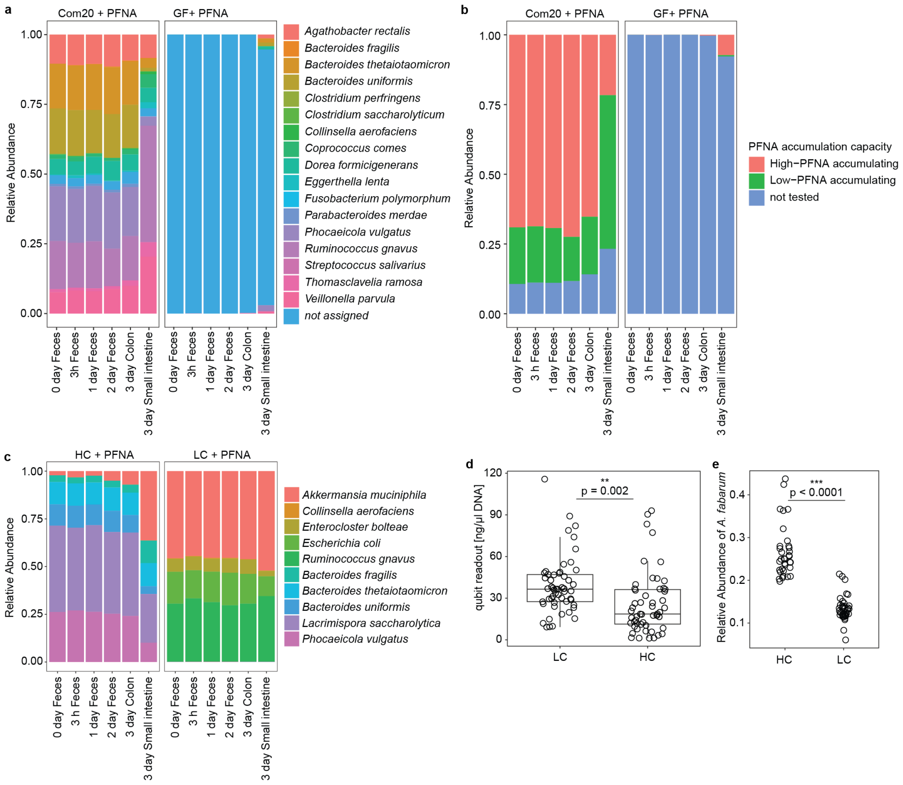


**Supplementary Figure 9. PFNA does not affect gut microbiota composition of mice colonised with human gut bacteria. a.** Gut microbiota composition of mice colonised with 20 human gut bacterial strains (Com20) and exposed to 10 mg/kg body weight by oral gavage on day 0. 17 out of 20 gut bacteria colonised the mice with 7 strains making up 90 % of bacterial composition (*P. vulgatus, R. gnavus, B. thetaiotaomicron, B. uniformis, A. rectalis, Veillonella parvula, Dorea formicigenerans*). The composition in the small intestine differed from that in the feces and colon, with the dominating species being *R. gnavus* and *V. parvula* (SI Table 41). **b.** In the colon and fecal samples 70 % of bacteria were classified as high-PFNA accumulating strains, while in the small intestine only 20 % of bacteria were classified as high-PFNA accumulating (SI Table 42). **c.** Gut microbiota composition of mice colonised with a community of five high-PFNA accumulating strains (HC) and five low-PFNA accumulating strains (LC) exposed to 10 mg/kg body weight by oral gavage on day 0. All gut bacteria colonised the mice, but *A. muciniphila* was present in both HC and LC colonised mice (SI Table 43). **d.** LC (low-PFNA accumulating community) colonised mice showed higher DNA yields after DNA extraction compared to HC (high-PFNA accumulating community) colonised mice (1.96-fold change, p-value = 0.002), indicating higher bacterial content in LC colonised mice (SI Table 39). **e.** HC colonised mice showed higher relative abundance of *Acetobacter fabarum* spike-in compared to LC colonised mice (2-fold change, p-value = 0.0009), indicating higher bacterial content in LC colonised mice. Subset of 76 samples (SI Table 44).

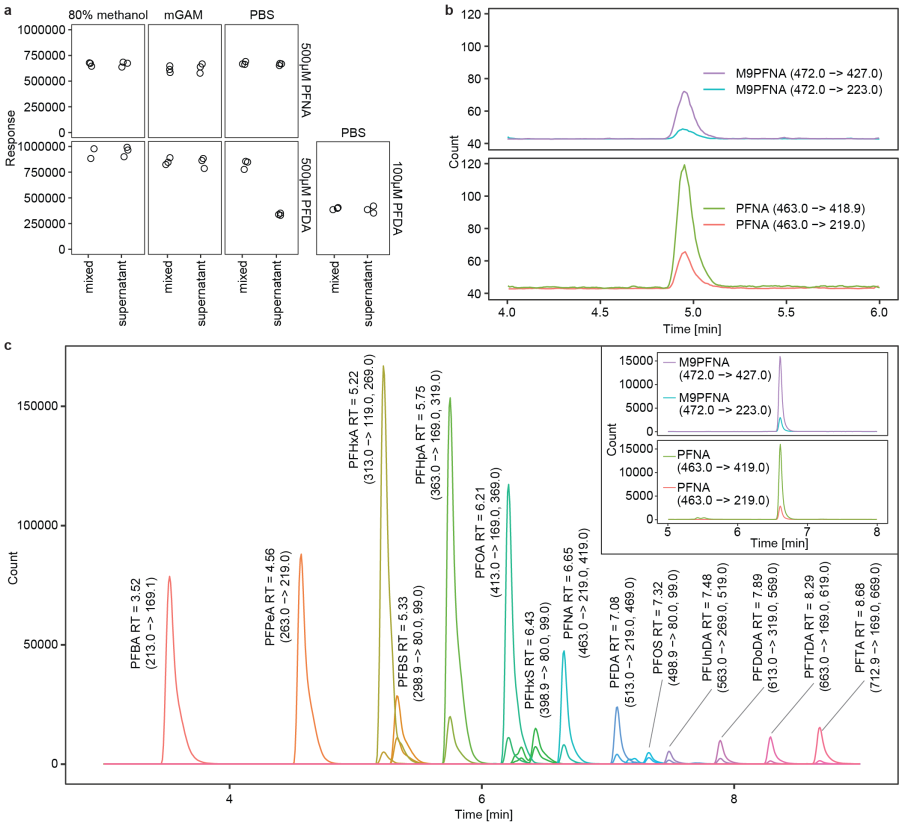

**Supplementary Figure 10. a.** Solubility of PFNA and PFDA in 80% methanol (1% DMSO), mGAM (1% DMSO) and PBS (1% DMSO). PFNA was soluble in all conditions up to 500 µM, while PFDA was soluble up to 500 µM in 80% methanol and mGAM, and up to 100 µM in PBS. n = 3 technical replicates (SI Table 45). **b.** PFNA and 13C-labelled PFNA chromatograms for 10 min reverse-phase LC method used with QQQ for mouse fecal sample analysis Fig. 5. **c.** Chromatograms for 14 PFAS compounds and their 13C labelled internal standards using the EPA Draft Method 1633 (Fig. 2d,f).
